# Vagal TRPV1^+^ sensory neurons regulate myeloid cell dynamics and protect against influenza virus infection

**DOI:** 10.1101/2024.08.21.609013

**Authors:** Daping Yang, Nicole Almanzar, Jingya Xia, Swalpa Udit, Stephen T. Yeung, Camille Khairallah, Daisy A. Hoagland, Benjamin D. Umans, Nicole Sarden, Ozge Erdogan, Nadia Baalbaki, Anna Beekmayer-Dhillon, Juhyun Lee, Kimberly A. Meerschaert, Stephen D. Liberles, Bryan G. Yipp, Ruth A. Franklin, Kamal M. Khanna, Pankaj Baral, Adam L. Haber, Isaac M. Chiu

## Abstract

Influenza viruses are a major global cause of morbidity and mortality. Vagal TRPV1^+^ nociceptive sensory neurons, which innervate the airways, are known to mediate defenses against harmful agents. However, their function in lung antiviral defenses remains unclear. Our study reveals that both systemic and vagal-specific ablation of TRPV1^+^ nociceptors reduced survival in mice infected with influenza A virus (IAV), despite no significant changes in viral burden or weight loss. Mice lacking nociceptors showed exacerbated lung pathology and elevated levels of pro-inflammatory cytokines. The increased mortality was not attributable to the loss of the TRPV1 ion channel or neuropeptides CGRP or substance P. Immune profiling through flow cytometry and single-cell RNA sequencing identified significant nociceptor deficiency-mediated changes in the lung immune landscape, including an expansion of neutrophils and monocyte-derived macrophages. Transcriptional analysis revealed impaired interferon signaling in these myeloid cells and an imbalance in distinct neutrophil sub-populations in the absence of nociceptors. Furthermore, anti-GR1-mediated depletion of myeloid cells during IAV infection significantly improved survival, underscoring a role of nociceptors in preventing pathogenic myeloid cell states that contribute to IAV-induced mortality.

**One Sentence Summary**: TRPV1^+^ neurons facilitate host survival from influenza A virus infection by controlling myeloid cell responses and immunopathology.

## INTRODUCTION

The respiratory tract is a crucial site for defense against various threats, including inhaled bacterial and viral pathogens. Influenza viruses are a leading cause of seasonal epidemics, accounting for millions of severe infections and more than a quarter million deaths annually worldwide (1, 2). Influenza A virus is particularly notable in causing epidemics and can be life-threatening to high-risk individuals, such as young children, the elderly, and those with conditions like chronic respiratory disease, obesity, and cardiovascular disease (1). A well-coordinated response to respiratory infection is essential for host survival, involving both the immune and nervous systems. While the immune system plays an integral role in antiviral resistance mechanisms and mediating viral clearance (3–5), an exaggerated inflammatory response can cause severe immunopathology and drive mortality during influenza infection (4, 6). Understanding the cellular mechanisms that mediate both respiratory antiviral defenses and resolution of inflammation could provide key insights into why certain individuals experience more severe disease and increased mortality risk.

The nervous system may play a critical role in coordinating host responses to pathogen infections and mediating the resolution of inflammation. The respiratory tract is densely innervated by sensory neurons that detect and respond to a diverse range of environmental signals, as well as by parasympathetic and sympathetic efferent neurons that regulate lung physiology (7–9). During infection, neural circuits are activated that drive sickness behaviors including malaise, loss of appetite, nasal congestion, headache, fever, sore throat, and cough (1, 3, 10). Sensory neurons can regulate pulmonary cytokine production and antimicrobial immunity by directly communicating with the immune system through the local release of neuropeptides and neurotransmitters (11, 12) or by indirectly activating brainstem circuits that elicit vagal efferent responses (13, 14).

Sensory innervation of the trachea and lungs is primarily provided by neurons of the vagus nerve, with their cell bodies located in the vagal nodose/jugular ganglia located at the base of the skull (7, 8, 15, 16). Among these neurons, a subset are classified as nociceptors due to their ability to detect and initiate responses to harmful stimuli such as chemical irritants, mechanical injury, and deviations in pH or temperature (7, 17). Nociceptors detect these stimuli via specialized receptors expressed at their nerve terminals, including members of the transient receptor potential (TRP) family of large-pore cation channels (18–20). A key molecular sensor of noxious stimuli expressed by nociceptors is TRPV1, a ligand-gated ion channel that detects capsaicin, protons, and high temperatures (18, 20, 21). Upon activation, airway-innervating vagal nociceptors signal to the brainstem, inducing reflexive responses such as bronchoconstriction and cough (7, 22).

Recent studies have revealed that nociceptors regulate immunity during allergic airway inflammation (23–25) and bacterial pneumonia (11, 26). This regulation occurs through the release of neuropeptides including calcitonin gene-related peptide (CGRP) and substance P (SP), which modulate innate and adaptive immune cell function in the airways (11, 24, 25). Vagal neurons have been found to change transcriptionally during influenza infections, and blockade of vagal neuron activity exacerbates infection outcomes in mice (27, 28). However, the functional role of TRPV1^+^ vagal nociceptors in modulating the immune response to influenza infection remains unclear.

In this study, we identify a critical role for TRPV1^+^ neurons in regulating myeloid cell-driven immunopathology in influenza A virus (IAV) infection. Using both chemical and genetic ablation strategies in mice, we demonstrate that vagal sensory neurons are essential for survival during IAV infection and for the proper regulation of proinflammatory cytokine induction, pulmonary macrophage responses, and neutrophil influx. Nociceptor-deficient mice exhibited reduced interferon signaling and imbalanced neutrophil populations, while myeloid cell ablation rescued these mice from IAV-induced mortality. Our findings highlight the role of the nervous system in preventing immunopathology and mediating host defense during viral lung infection.

## RESULTS

### Nociceptive sensory neurons are crucial for survival during Influenza A virus infection

The airways are innervated by sensory neurons that play distinct roles in lung function and physiology (7, 8, 29, 30). Nociceptors that reside in the airways functionally express ion channels including TRPV1 and Nav1.8 (12, 20). TRPV1 is a ligand-gated cation channel that detects capsaicin, protons, and noxious heat (20). Nav1.8 is a voltage-gated sodium channel that mediates action potential generation in sensory neurons, labeling a large proportion of nociceptors (12, 20).

We first visualized lung innervation by TRPV1^+^ nociceptors using *Trpv1*-Cre mice injected neonatally with Adenovirus-Associated Virus (AAV9) that expresses tdTomato under a Cre-dependent promoter. Imaging of adult mouse lung sections showed a network of nerves surrounding large and small airways labeled with pan-neuronal marker Tuj1; a subset of these neurons were TRPV1-tdTomato^+^ (**Fig. 1A**). To explore the role of TRPV1^+^ sensory neurons in IAV infection pathogenesis, we used a well-established method for chemical ablation of nociceptors. Wildtype B6 mice were pretreated with resiniferatoxin (RTX), a potent TRPV1 agonist that induces excitotoxicity-mediated cell death of TRPV1^+^ neurons (11, 31, 32). Four weeks after RTX treatment, the mice were intranasally inoculated with 50 TCID_50_ of the H1N1 IAV strain PR8 (A/Puerto Rico/8/1934). RTX-treated mice exhibited a significantly lower survival rate compared to vehicle-treated controls (**Fig. 1B**). This increased mortality was observed in both male and female mice (**Fig. 1B**).

**Figure 1.**
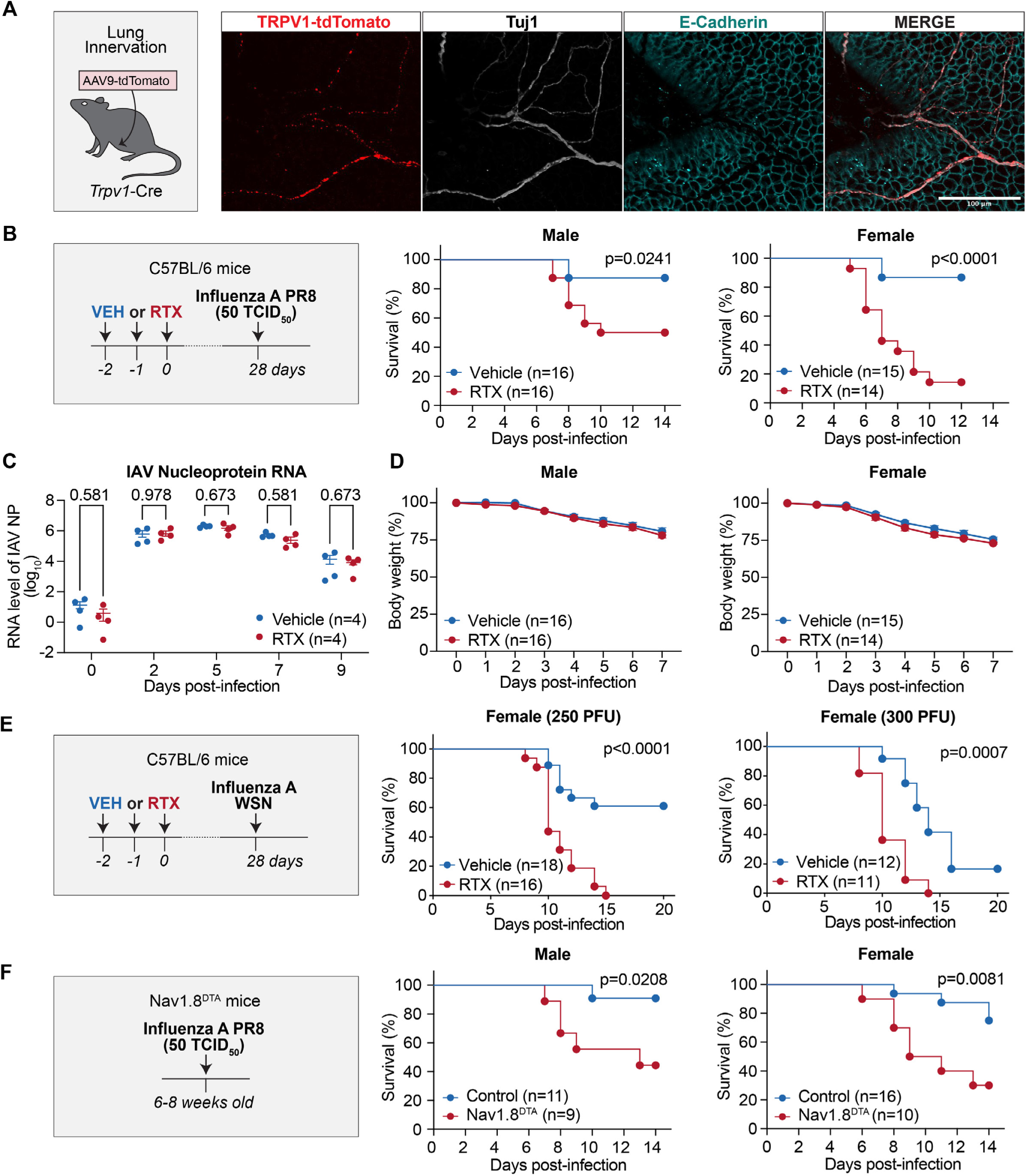
Nociceptors promote host survival against influenza infection. **(A)** Maximum intensity projection image of lung tissue from *Trpv1*-Cre mice injected at post-natal day 1 with AAV9-tdTomato (schematic) and immunostained for Tuj1 and E-cadherin. Scale bar 100µm. **(B)** C57BL/6 mice were treated with RTX to chemically ablate TRPV1^+^ neurons then intranasally infected with the influenza A virus (IAV) PR8 (50 TCID_50_ per mouse) four weeks later (schematic). Survival curves shown for male (left, n=16 per group) and female (right, n=14-15 per group) vehicle and RTX-treated mice. **(C)** IAV nucleoprotein (NP) transcript levels were measured in the lungs from infected mice at indicated time points post-infection in RTX vs. vehicle treated mice (n=4 mice per group). **(D)** Change in body weight as a percentage of initial body weight (y axis) of male (left, n = 16 per group) and female (right, n=14-15 per group) vehicle and RTX-treated mice post-infection with IAV PR8. **(E)** C57BL/6 mice were treated with RTX then intranasally infected with IAV WSN four weeks later (schematic). Survival curves shown for female vehicle and RTX-treated mice infected with 250 PFU (left, n=16-18 per group) or 300 PFU (left, n=11-12 per group) of IAV WSN. **(F)** Survival curves of Nav1.8^DTA^ mice and littermate controls intranasally infected with IAV PR8 (50 TCID_50_ per mouse, schematic), shown separately for male (left, n = 9-11 per group) and female (right, n=10-16 per group) mice. Log-rank test was used in (B, E-F); Multiple t test in (C and D); error bars show mean±SEM; P values labeled.

We next investigated whether nociceptors influenced viral clearance by measuring IAV nucleoprotein (NP) mRNA levels (33). Following PR8 infection, viral NP transcript levels increased significantly in lung tissues, as expected; however, there were no differences between RTX- and vehicle-treated mice at any time point during the 9 days of infection (**Fig. 1C**). Previous studies have shown that metabolic intake affects the outcome of viral infections (34), and that *Gabra1*^+^ sensory neurons regulate food consumption during influenza infections (10). Therefore, we examined whether ablation of TRPV1^+^ sensory neurons affected weight loss or food consumption. We found no differences in food or water intake between RTX- and vehicle-treated mice (**Fig. S1A**). Additionally, weight loss up to the point of mortality was comparable between male and female RTX-treated mice and vehicle controls (**Fig. 1D**).

To test whether the protection offered by TRPV1^+^ neurons was specific to the IAV strain PR8, we infected mice with H1N1 IAV strain WSN (A/WSN/1933). As with PR8 infection, we found that RTX-treated mice showed worsened survival from two doses (250 PFU and 300 PFU) of IAV WSN infection compared to vehicle-treated mice (**Fig. 1E**). All together, these data demonstrate a critical role for TRPV1^+^ nociceptors in mediating survival against IAV infection that is independent of viral clearance and weight loss.

As a second strategy to ablate nociceptors, we utilized a genetic approach to target Nav1.8-lineage neurons (12, 20). Nav1.8-Cre mice were crossed with Cre-dependent Diptheria Toxin A (DTA) reporter mice to generate mice lacking Nav1.8^+^ nociceptors (Nav1.8^DTA^ mice) and Cre-negative control littermates. Following IAV PR8 intranasal inoculation, Nav1.8^DTA^ mice showed significantly decreased survival compared to control mice, in both male and female mice (**Fig. 1F**). To better understand how nociceptor ablation altered the physiologic response to IAV infection, we assessed vital signs in naïve mice and at 7 days post-infection (DPI), a time point where survival began to diverge in RTX vs. vehicle-treated mice (**Fig. 1B**) and Nav1.8^DTA^ vs. control littermates (**Fig. 1F**). At baseline, RTX-treated and Nav1.8^DTA^ mice showed no difference in oxygen saturation, heart rate, or breathing rate (**Fig. S1B-C**). The percent oxygen saturation decreased after infection in all mice, but both RTX-treated and Nav1.8^DTA^ mice exhibited more marked reductions with lower oxygen saturation at 7 DPI compared to their respective controls (**Fig. S1B-C**). In vehicle-treated mice (no RTX), IAV infection caused a drop in heart rate compared to baseline whereas heart rate remained unchanged in RTX-treated mice; Nav1.8^DTA^ mice and controls both exhibited decreased heart rate upon IAV infection (**Fig. S1B-C**). Breathing rate did not change after IAV infection and showed no difference in nociceptor-deficient mice compared to controls (**Fig. S1B-C**).

### TRPV1 ion channel does not affect influenza infection outcome

While TRPV1 is sensitized during inflammation to drive heat sensitivity and mediates detection of noxious stimuli (20), its role in viral infection is unknown. We next performed PR8 infections in TRPV1-deficient mice (*Trpv1*^−/−^) and wild-type controls, finding no difference in mortality (**Fig. S2A**) or bodyweight loss following infection (**Fig. S2B**). *Trpv1*^−/−^ mice also exhibited similar oxygen saturation levels, heart rates, and breathing rates compared to controls at 7 DPI (**Fig. S2C**). Therefore, while TRPV1-expressing neurons are required to mediate host protection, the survival defects and differences in vital signs after IAV infection are not mediated by the TRPV1 ion channel.

### Vagal TRPV1^+^ neurons upregulate ATF3 and mediate survival during influenza infection

The sensory innervation of the lungs is primarily provided by the vagus nerve (7, 8). We confirmed significant expression of TRPV1 in vagal nodose and jugular sensory ganglia through immunostaining (**Fig. 2A**). To visualize the vagal TRPV1^+^ innervation of the lungs, we injected AAV9 carrying a Cre-dependent tdTomato reporter into the vagal ganglia of *Trpv1*-Cre mice (**Fig. 2B**). This approach revealed anterogradely traced vagal TRPV1^+^ nerve fibers around the lung airways (**Fig. 2B**). We hypothesized that vagal nociceptors respond to IAV infection by upregulating the transcription factor ATF3, an adaptive response gene activated by neuronal injury and other stressors (35, 36). Immunostaining showed increased ATF3 positivity in vagal sensory neurons post-IAV infection, with approximately 20% of total neurons and 40% of TRPV1^+^ neurons staining positive for ATF3 at 7 DPI (**Fig. 2D**). This finding supports previous studies indicating significant transcriptional changes in vagal sensory neurons following influenza infection (27, 28).

**Figure 2.**
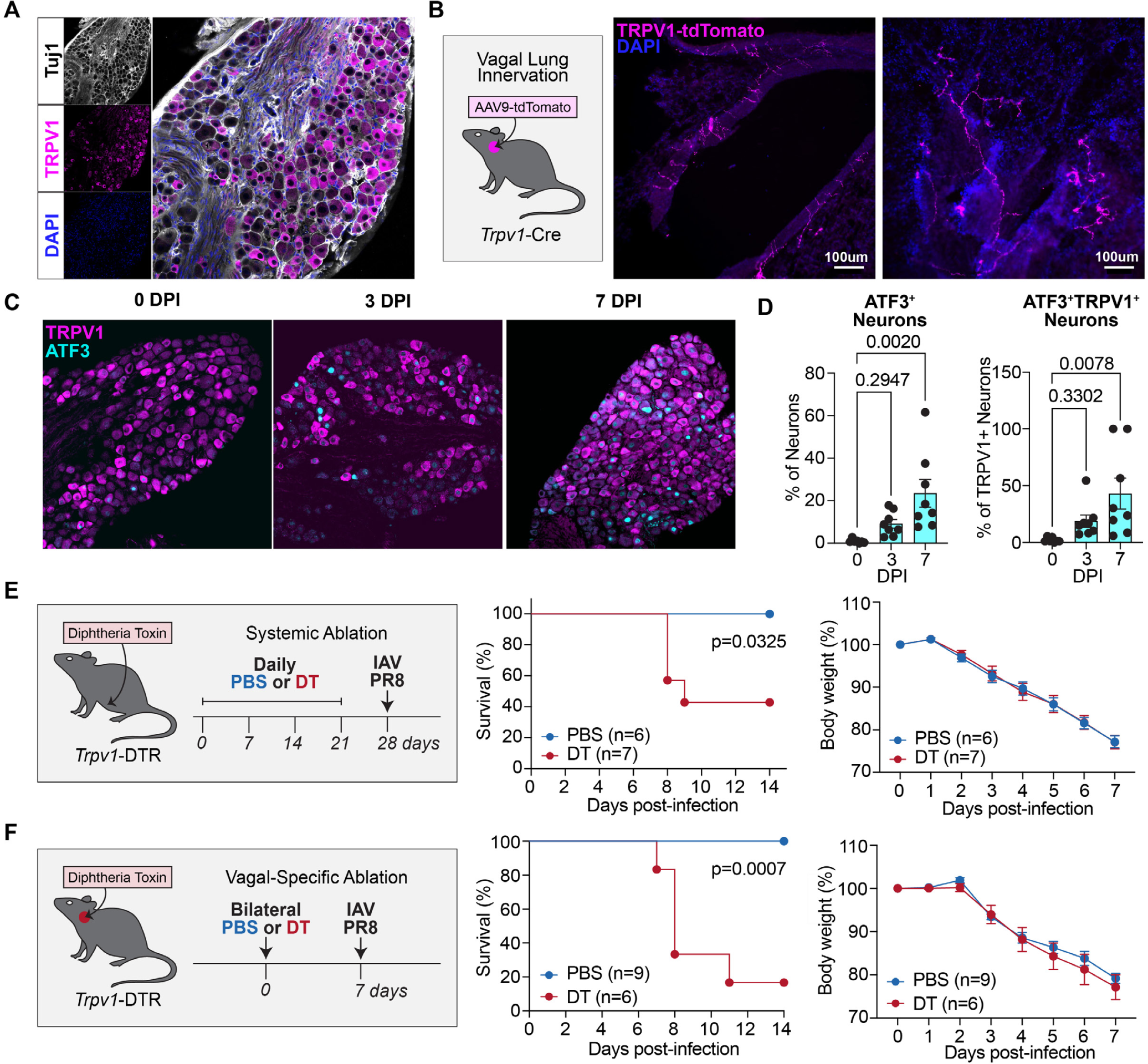
Vagal TRPV1+ sensory neurons mediate host protection against influenza infection. **(A)** Immunostaining of TRPV1, Tuj1 and DAPI in the jugular-nodose ganglia collected from wild-type mice. **(B)** Images of vagal *Trpv1*^+^ nerve fibers labeled in the lung tissue of *Trpv1*-Cre mice injected with AAV9-FLEX-tdTomato into the jugular-nodose ganglia (schematic) and counterstained with DAPI. **(C)** Representative images of jugular-nodose ganglia from wild-type mice at indicated timepoints post-IAV PR8 infection, showing immunostaining for TRPV1 and ATF3. **(D)** Quantification of the percentage of all neurons (left) and percentage of TRPV1^+^ neurons (right) positive for ATF3 per ganglia. **(E)** *Trpv1*-DTR mice received daily intraperitoneal injections of diphtheria toxin (DT, 200 ng per mouse) for three weeks to systemically ablate *Trpv1*^+^ neurons and were then intranasally infected with IAV PR8 a week later (schematic). Survival curves (left) and change in body weight (right) were monitored post-infection (n = 6-7 per group). **(F)** *Trpv1*-DTR mice received bilateral injections of DT (20 ng per mouse) into the jugular-nodose ganglia to specifically ablate *Trpv1*^+^ vagal neurons and then infected with IAV PR8 a week later (schematic). Survival curves (left) and change in body weight were monitored post-infection (n= 6-9 per group). One-way ANOVA in (D and F); Log-rank test was used for survival analysis; Multiple t test for body weight analysis; error bars show mean±SEM; P values labeled.

To investigate the role of TRPV1^+^ vagal nociceptors in IAV infection specifically, we used *Trpv1*-DTR mice, which express the diphtheria toxin receptor (DTR) under the control of the *Trpv1* promoter, allowing targeted ablation of *Trpv1*^+^ neurons via diphtheria toxin (DT) injection (11, 37). Systemic DT administration delivered via peritoneal injection in *Trpv1*-DTR mice, which results in ablation of all *Trpv1*^+^ neurons, recapitulated the reduced survival observed in RTX-treated and Nav1.8-DTA mice, without impacting body weight loss (**Fig. 2E**). For vagal specific ablation, we performed bilateral intra-ganglionic DT injections into the nodose/jugular ganglia of *Trpv1*-DTR mice and allowed to recover for a week before infecting them with IAV PR8. Vagal-specific ablation of nociceptors led to a significant reduction in survival compared to vehicle-injected controls, with no difference in body weight loss (**Fig. 2F**). Therefore, vagal TRPV1^+^ nociceptors are crucial for host protection against IAV infection.

### Influenza infection outcome is not mediated by neuropeptides CGRP or substance P

After identifying nociceptors as crucial for survival in influenza infection, we explored the mechanisms behind this host defense. Re-analyzing a single-cell RNA sequencing dataset of vagal sensory neurons (GEO: GSE145216), we confirmed that *Trpv1* is expressed in a large subset of these neurons (**Fig. S3A**) (29). Most *Trpv1*^+^ neurons (66%) expressed *Scn10a* (Nav1.8), and also showed co-expression with either *Prdm12* (a marker for jugular neurons) or *Phox2b* (a marker for nodose neurons), indicating their presence in both the jugular and nodose ganglia (**Fig. S3C-D**). Neuropeptides CGRP and substance P, encoded by *Calca* and *Tac1* respectively, were highly expressed in *Trpv1*^+^ jugular neurons, with some presence in nodose clusters (**Fig. S3C-D**). As previously described (10, 29, 30), *Trpv1*^+^ neurons overlapped with *Npy2r*^+^ and *Npy1r*^+^ populations and were distinct from *P2ry1*^+^ populations (**Fig. S3D**). *Gabra1*^+^ neurons, recently implicated in driving sickness behaviors and weight loss during influenza virus infections (10) were distinct from the *Trpv1*^+^ vagal neurons (**Fig. S3D**).

Given the high level of expression of the neuropeptides CGRP and substance P by TRPV1^+^ vagal neurons, we tested their contribution to host defense in IAV infection. After PR8 infection, *Calca*^−/−^ mice, which lack CGRP, and wild-type controls exhibited no differences in survival or body weight loss at both low and high IAV inoculation doses (**Fig. S4A-B**). *Tac1*^−/−^ mice, which lack substance P, also demonstrated no differences in survival or body weight loss compared to wild-type controls at two infectious doses (**Fig. S4C-D**). These data suggest that the protective role of nociceptive sensory neurons in IAV infection does not depend on the neuropeptides CGRP or substance P.

### Nociceptor-deficient mice display exacerbated lung pathology and increased flu-induced cytokine levels

To investigate why nociceptor ablation leads to reduced survival, we conducted a histopathological analysis of lung tissue from naïve uninfected mice and at 7 DPI. At baseline, the lungs of RTX-treated mice appeared similar to those of vehicle-treated controls (**Fig. 3A**). However, at 7 DPI, RTX-treated mice displayed heightened alveolar hemorrhage, diffuse alveolar damage, immune cell infiltration, and more severe pulmonary edema, particularly when examined via non-perfused lung sections (**Fig. 3B**, bottom row). Additionally, RTX-treated mice had detectable blood in the bronchoalveolar lavage fluid (BALF), which was absent in vehicle-treated controls (**Fig. S5A**), consistent with the pathological finding of alveolar hemorrhage. Histopathological scoring by blinded observers confirmed increased immune infiltrates and greater pathological damage in lung sections from RTX-treated mice (**Fig. 3C**).

**Figure 3.**
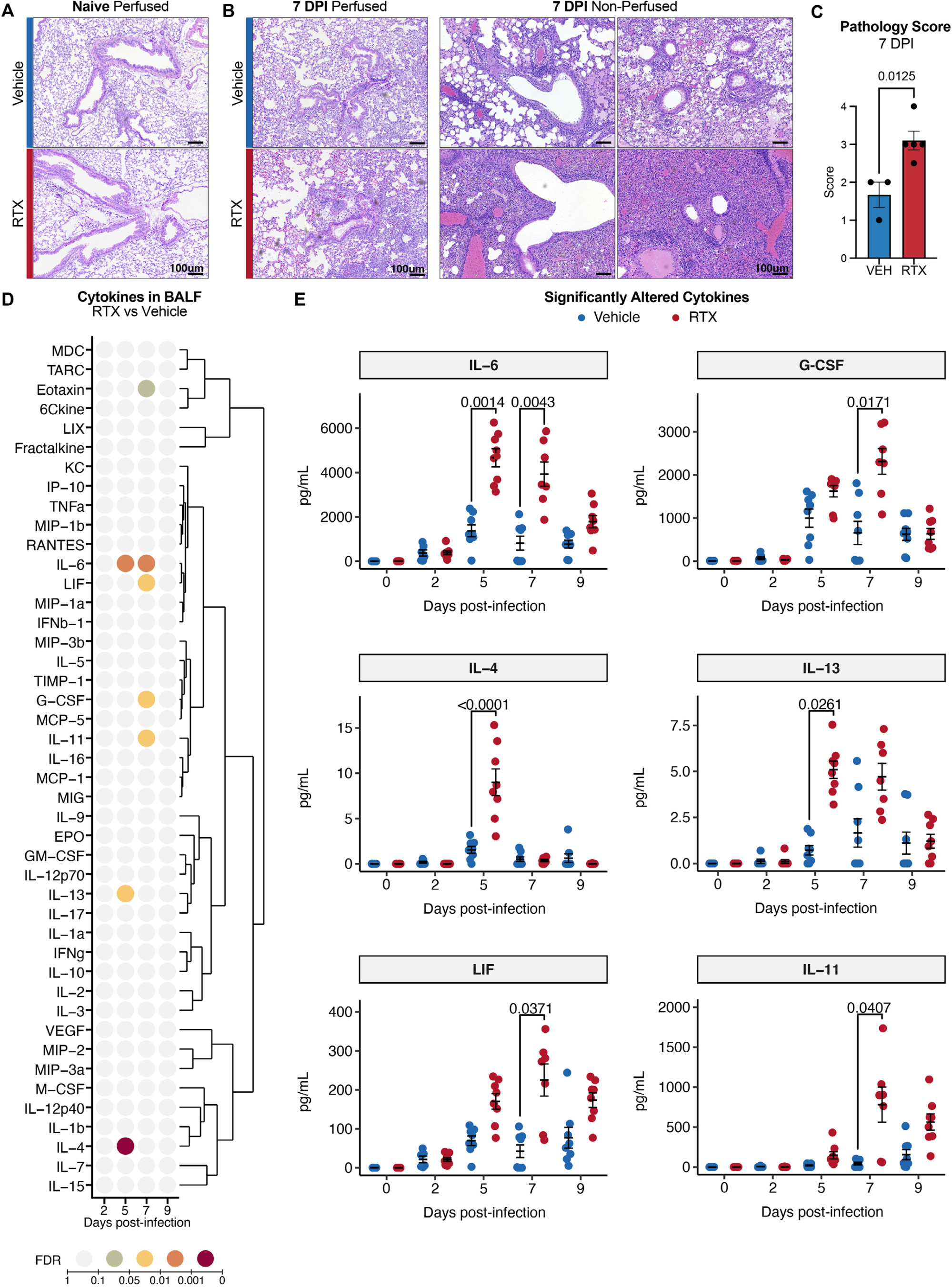
Nociceptor ablated mice displayed enhanced pulmonary pathology and altered cytokine induction after influenza infection. **(A)** Representative H&E images of perfused lung tissue from uninfected vehicle (top) and RTX-treated (bottom) mice. **(B)** Representative H&E images of perfused (left) and non-perfused (right) lung tissue from vehicle (top) and RTX-treated (bottom) mice at 7 days post IAV PR8 infection. **(C)** Scoring of lung pathology observed in non-perfused H&E images at 7 days post-infection. **(D)** Protein levels of indicated cytokines (y axis) were detected in BALF at various timepoints (x axis) after IAV PR8 infection and measured using the 44-Plex Discovery Assay® Array (n = 4-8 per group). Color of dots illustrates significance of FDR-adjusted p value (color legend) for the comparison between RTX and vehicle of each cytokine at each time point after infection. Cytokines were hierarchically clustered (dendrogram) according to their detected protein levels. **(E)** Detected protein levels (in pg/mL, y axis) of the six cytokines significantly different between vehicle and RTX, measured in BALF at days 0 (naïve), 2, 5, 7 and 9 post-infection (x axis). Points colored by treatment group. (n = 4-8 per group). Scale bars 100um in (A-B); Student’s t test in (C); Two-way ANOVA with FDR adjusted p values in (D-E); P values labeled.

We hypothesized that loss of nociceptors might alter the inflammatory response to IAV infection. To test this, we employed multiplex ELISAs to measure levels of 44 cytokines in BALF collected from naïve mice and from RTX and vehicle-treated mice at days 2, 5, 7, and 9 post-infection (**Fig. 3D** and **Fig. S5B**, **Table S1**). After correcting for multiple testing, six cytokines exhibited significant differences (FDR < 0.05) between RTX and vehicle-treated mice (**Fig. 3D-E**). IL-6, a cytokine crucial for antiviral defense (38, 39) but also linked to lung immunopathology (40, 41), was significantly elevated in RTX-treated mice at days 5 and 7 post-infection (**Fig. 3D-E**). G-CSF, a cytokine that stimulates differentiation of bone marrow precursor cells into neutrophils and monocytes/macrophages (42) and is associated with severe viral pneumonia (40), was elevated in RTX-treated mice at day 7 post-infection (**Fig. 3D-E**). RTX-treated mice also exhibited increased levels of type 2 cytokines IL-4 and IL-13, and higher levels of the IL-6 family cytokine LIF and the fibrosis-inducing cytokine IL-11 (43) compared to vehicle-treated mice at day 5 post-infection (**Fig. 3D-E**).

We further complemented the cytokine array by measuring levels of IFNα and IFNλ with additional ELISAs (**Fig. S6A-B**). In vehicle-treated mice, IFNα levels peaked early after infection and declined by day 7, whereas RTX-treated mice showed only a modest increase starting at day 5, indicating altered IFNα dynamics (**Fig. S6A**). No significant difference in IFNλ levels was observed between RTX and vehicle-treated mice (**Fig. S6B**). In summary, we observed a robust induction of several proinflammatory cytokines (IL-6, G-CSF, IL-4, IL-13, LIF and IL-11) that was significantly higher in nociceptor deficient mice compared to control animals.

### Nociceptors regulate monocyte and neutrophil recruitment during influenza infection

After observing that nociceptor-deficient mice showed signs of increased inflammation by lung histology and elevated cytokine levels in BALF, we quantified changes in immune cells with flow cytometry. The host immune response to respiratory viral infections requires both the rapid response of lung-resident myeloid cells and lymphocytes, as well as the recruitment of infiltrating myelomonocytic cells; these components can both mediate antiviral immunity and drive immunopathology (3). We measured myeloid immune cells (**Fig. 4A**) and lymphocytes (**Fig. S9A**) in lung tissues from vehicle- and RTX-treated mice at baseline and 7 DPI. We found that the number of neutrophils – gated as CD11c^−^CD11b^+^Ly6G^+^ cells (**Fig. 4A**) given the appearance of Ly6G^+^ monocyte-derived macrophages in the lungs after influenza infection (44) – significantly increased in RTX-treated mice at 7 DPI compared to vehicle-treated mice, while there was no significant difference in neutrophils at baseline (**Fig. 4B**). After infection, we observed significant increases in lung monocytes (CD11b^+^F4/80^+^Ly6C^Hi^CD11c^Lo^) (**Fig. 4C**) and monocyte-derived macrophages (CD11b^+^F4/80^+^Ly6C^Hi^CD11c^Hi^) (**Fig. 4D**). At 7 DPI, monocyte-derived macrophages were strikingly elevated in RTX-treated mice compared to vehicle-treated controls both as a fraction of immune cells and in absolute cell numbers (**Fig. 4D**). A population of CD11b^+^F4/80^+^Ly6C^Lo^CD11c^Lo^ cells, which likely consists of both Ly6C^Lo^ monocytes and lung interstitial macrophages, also showed an increase in RTX-treated mice at 7 DPI (**Fig. 4E**). By contrast, alveolar macrophages (CD11c^+^SiglecF^+^) showed a slight decrease with infection but were unchanged between RTX and vehicle-treated mice (**Fig. S8A**). We also collected BALF at 7 DPI and found a significant increase in both neutrophils and monocytes in RTX-treated mice compared to vehicle-treated mice (**Fig. S7A-B**). These findings suggest that the loss of nociceptors leads to dysregulated recruitment of neutrophils and monocytes, and an expansion of monocyte-derived macrophages.

**Figure 4.**
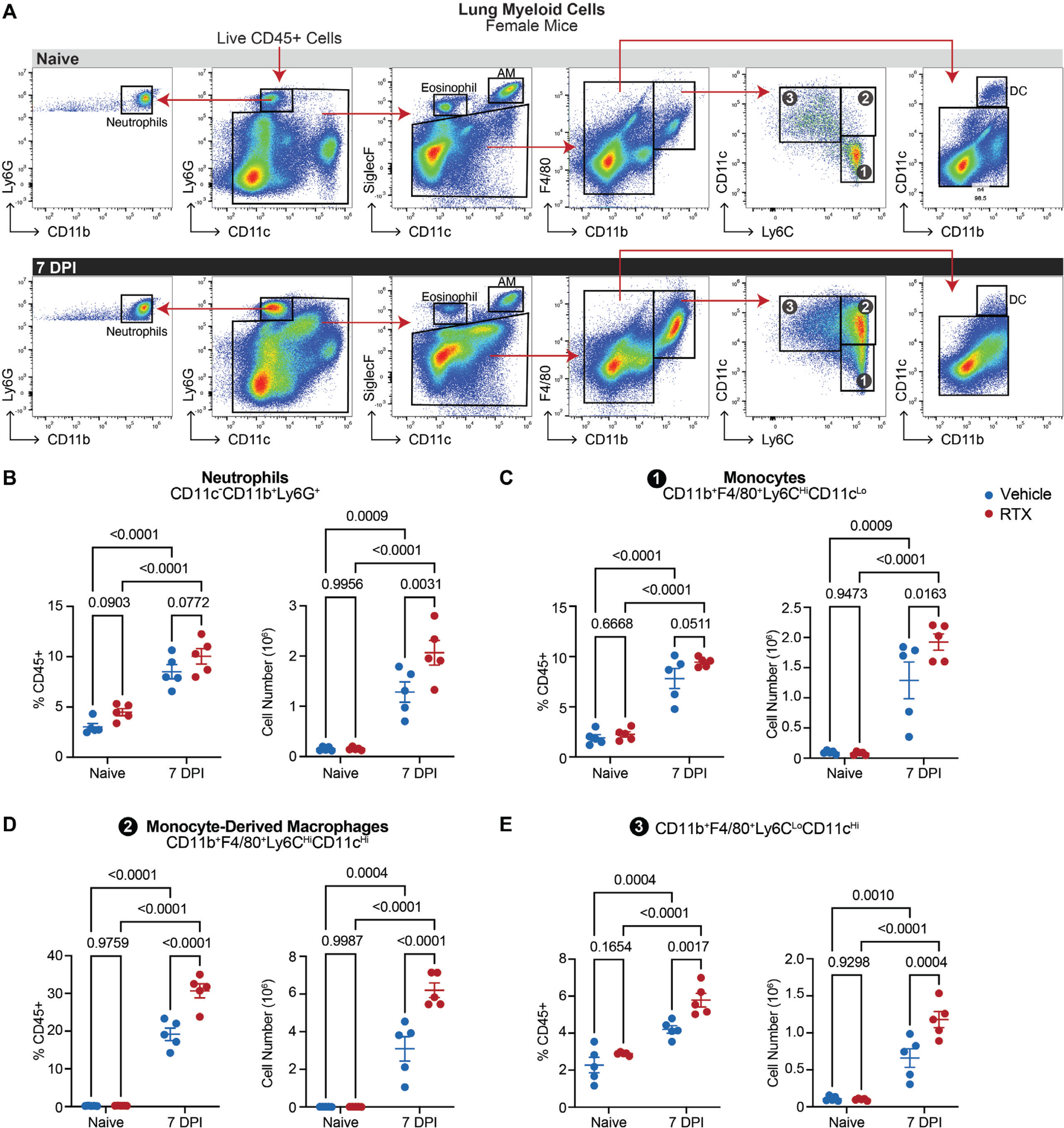
Influenza-induced myeloid cell responses are dysregulated in nociceptor deficient mice. **(A)** Representative flow cytometry plots and gating strategy for the labeled myeloid immune cell populations isolated from the lungs of naïve mice (top) and at day 7 post IAV PR8 infection (bottom). **(B-E)** Quantification of lung myeloid cell populations from vehicle- and RTX-treated female mice at baseline and at 7 DPI as a percentage of CD45^+^ cells (left) or absolute cell numbers (right), shown for **(B)** neutrophils (CD11c^−^CD11b^+^Ly6G^+^), **(C)** monocytes (CD11b^+^F4/80^+^Ly6C^Hi^CD11c^Lo^), **(D)** monocyte-derived macrophages (CD11b^+^F4/80^+^Ly6C^Hi^CD11c^Hi^), and (**E**) CD11b^+^F4/80^+^Ly6C^Lo^CD11c^Hi^ cells. (n=5 per group). Two-way ANOVA in (B-E); error bars show mean±SEM; P values labeled.

Eosinophils (CD11c^−^SiglecF^+^) and dendritic cells (CD11b^Lo^CD11c^+^) were a small proportion of the lung immune cells both at baseline and after infection and showed little difference between RTX and vehicle-treated mice (**Fig. S8A-B**). The differences between myeloid populations in RTX compared to vehicle-treated mice at 7 DPI were largely reproduced in a second independent experiment conducted in male mice (**Fig. S8D**). Our in-depth characterization of lung lymphoid cells did not reveal any differences in lymphocyte populations between RTX- and vehicle-treated mice at 7 DPI (**Fig. S9B**). Collectively, this data demonstrates that RTX treatment increases the number of neutrophils, monocytes, and monocyte-derived macrophages in the lungs at 7 DPI, implicating these cells as potential drivers of pathogenic inflammation in RTX-treated mice after IAV infection.

### Nociceptors do not regulate influenza-specific CD8^+^ T cell responses

Cytotoxic CD8^+^ T cells are crucial for protection against influenza by killing infected cells (45), and influenza infection leads to activation and expansion of virus-specific CD8^+^ T cells that can be detected by MHC-class I tetramer staining for the viral nucleoprotein (NP) (46). Of note, infection studies above were performed at Harvard and Yale University, and the following experiments were performed by coauthors at NYU. Consistent with previous experiments, we found that PR8 influenza infection in RTX-treated mice led to significantly worsened survival compared to vehicle-treated controls at both higher (75 EID_50_) and lower titers (50 EID_50_) of inoculum (**Fig. S10A-B**). Weight loss also did not differ between groups (**Fig. S10A-B**). To assess whether nociceptors modulate induction of virus-specific CD8^+^ T cells, we stained for NP-tetramer^+^ CD8^+^ T cells over the course of influenza infection (**Fig. S10**). We found no significant differences in CD4^+^ or CD8^+^ T cell numbers over time (**Fig. S10C-D**), and NP-tetramer^+^ CD8^+^ T cells between RTX-treated mice and control mice (**Fig. S10F-G**). These data indicate that nociceptors do not regulate T cell recruitment or induction of IAV-specific T cell responses.

### Characterization of lung immune dysregulation in nociceptor-deficient mice

To further define the role of nociceptors in regulating the immune compartment of the lung during IAV infection, we performed single-cell RNA sequencing (scRNAseq) after RTX-induced ablation of nociceptors. Briefly, whole lungs were isolated from vehicle and RTX-treated naïve animals or IAV-infected animals at 7 DPI and dissociated into a single-cell suspension that was then enriched for CD45^+^ immune cells before processing for scRNAseq (**Fig. 5A**). We performed unsupervised clustering and annotated cell types based on expression of known marker genes (**Fig. S11A-D**, **Table S2**). We identified low quality cells and doublets using technical quality metrics (Methods); these were retained in the dataset for visualization purposes but excluded from downstream analyses. Subsequently, we defined cell type-specific markers for each cluster in the dataset, which contained known lung myeloid and lymphoid immune cell types (**Fig. S11B**, **Table S3**).

**Figure 5.**
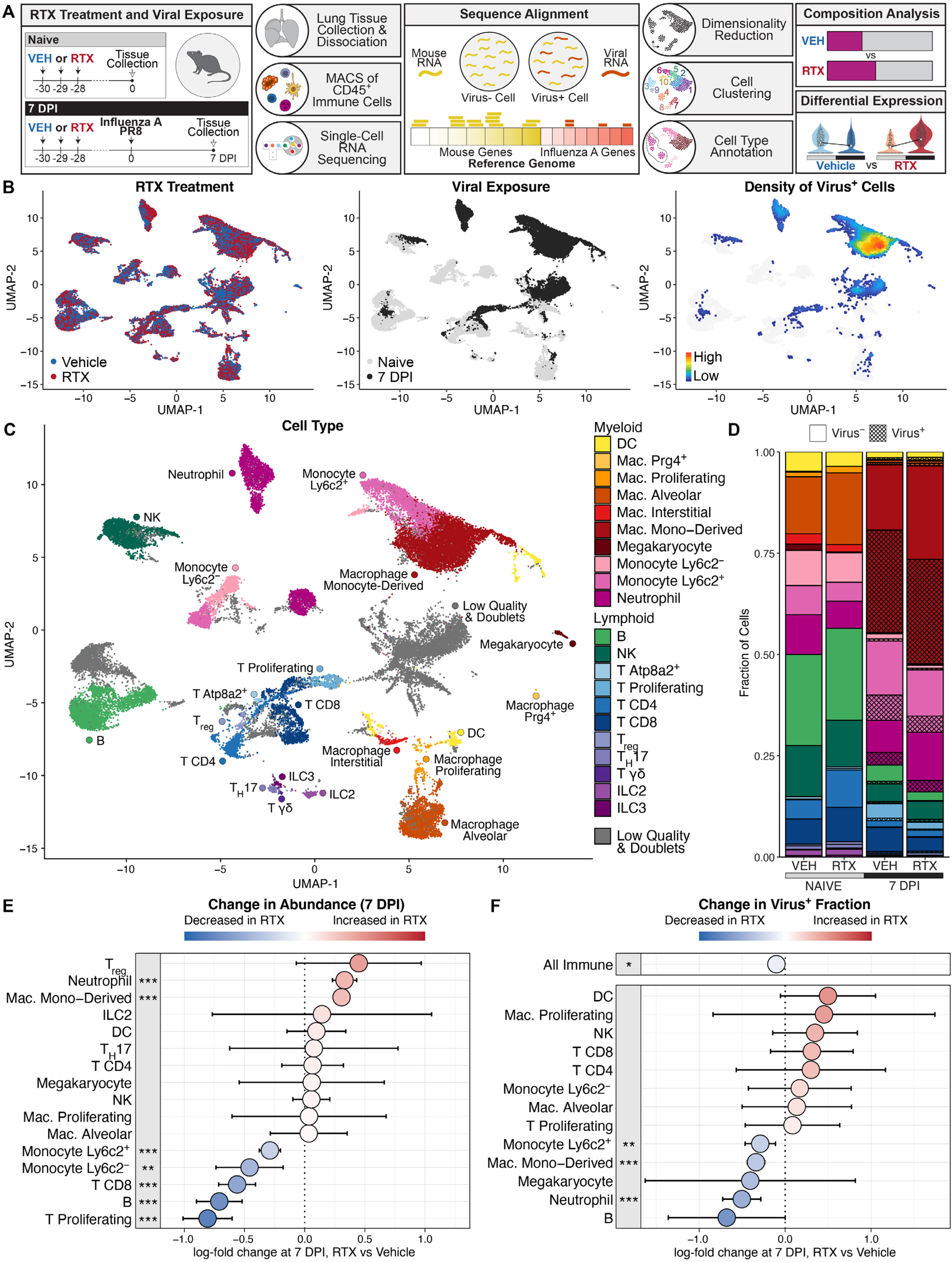
Nociceptor ablation alters the lung immune landscape following influenza infection. **(A)** Schematic representation of the experimental timeline, tissue processing workflow, and computational pipeline for scRNAseq dataset collection and analysis. **(B)** Uniform manifold approximation and projection (UMAP) of 37,543 lung immune cells (n = 2-3 mice pooled for each of the four groups) colored by treatment condition (left), viral exposure (middle), and density of Virus^+^ cells (right). **(C)** UMAP of cells colored and labeled by assigned cell type annotations (color legend and labels). **(D)** Cumulative bar plot indicating the fraction of immune cells (y axis) for each group (x axis) that are of each type (color legend on left); cross-hatch pattern (legend on top) overlayed to indicate cells of each type that contained viral transcripts. **(E)** Estimated binomial regression coefficient (Material and Methods) modeling the log-fold change in abundance (x axis, color legend) of each cell type (y axis). Negative and positive log-fold changes correspond to a decrease and increase, respectively, in RTX compared to vehicle at 7 days post-infection; colored dot: point estimate; bar: 95% confidence interval; asterisks indicate adjusted p value: ***: < 0.001, **: <0.01, *: <0.05. **(F)** Estimated binomial regression coefficient modeling the log fold change in Virus^+^ fraction (x axis, color legend) out of all immune cells (top) or of each cell type (bottom) (y axis) in RTX compared to vehicle at day 7 post-infection, visualized as in (E).

Visualization using the Uniform Manifold Approximation and Projection (UMAP) algorithm showed that cells from vehicle- and RTX-treated animals were distributed across clusters, whereas cells from naïve and 7 DPI primarily clustered apart from one another, indicating a profound effect of viral infection on the immune cell transcriptome, as expected (**Fig. 5B**). scRNAseq reads were aligned to the influenza viral genome which confirmed that IAV mRNA was detected (Methods) in a subset of cells—all of which came from the 7 DPI timepoint, as expected—that we refer to as Virus^+^ cells. Virus^+^ cells were most densely distributed within myeloid clusters, particularly monocyte-derived macrophages (**Fig. 5B-C**).

At baseline, myeloid cells and lymphocytes were detected in similar proportions in the lungs of both vehicle- and RTX-treated mice, with predominantly alveolar macrophages and B cells in each compartment, respectively (**Fig. 5D**). After IAV infection, we observed a strong expansion of the myeloid compartment primarily driven by the appearance of monocyte-derived macrophages, absent at baseline (**Fig. 5D** and **Fig. S12A**). Testing for changes in cell type composition, we found that at 7 DPI, RTX-treated mice had significantly more neutrophils and monocyte-derived macrophages compared to vehicle-treated mice, and fewer monocytes, CD8^+^ and proliferating T cells, and B cells (**Fig. 5E**). Viral transcripts were detected in fewer immune cells in RTX-treated mice; *Ly6c2*^+^ monocytes, monocyte-derived macrophages, and neutrophils all showed significant reductions in Virus^+^ cells (**Fig. 5F**). Given the recognized phagocytic roles of these cell types, this may be explained by an expansion of myeloid cells in RTX-treated mice relative to the viral burden, resulting in relatively more Virus^−^ ‘bystander’ cells. Together, this data demonstrates that nociceptor ablation altered the composition of the lung immune compartment after IAV infection and resulted in reduced frequency of Virus^+^ immune cells.

### Nociceptor ablation blunts influenza-induced interferon signaling in lung immune cells

To identify direct differences between vehicle and RTX-treated mice, we first performed differential expression comparisons between treatment groups at baseline and at 7 DPI (**Table S4**). At baseline, interstitial macrophages had the greatest number of genes differentially expressed (**Fig. S12B**), including an upregulation of heat shock proteins *Hspa1a* and *Hspa1b* and the tyrosine kinase *Mertk* in RTX (**Fig. S12D**). After influenza infection, neutrophils and *Ly6c2*^+^ monocytes showed the highest number of differentially expressed genes (**Fig. S12C**) with both cell types demonstrating downregulation of interferon-associated genes such as *Isg15* and *Cxcl10* in RTX (**Fig. S12E**).

To understand how RTX treatment affects the transcriptional response of each cell type to infection, we used a linear regression model to test for two-way interactions between RTX treatment and influenza exposure. For each cell type, this model identified genes that exhibit a response to infection that is significantly modified by RTX treatment, which we refer to as *differentially regulated* (**Fig. 6A**, **Table S4**). Myeloid cells were the most perturbed: neutrophils (214 genes), *Ly6c2*^+^ monocytes (204 genes), and alveolar macrophages (145 genes) (**Fig. 6B**). Given that these cells were also highly abundant, we normalized for cell number and found that neutrophils and *Ly6c2*^+^ monocytes remained the most transcriptionally altered after adjustment, suggesting that RTX treatment had a greater effect on the transcriptional response to infection in these two cell types (**Fig. 6C**). Pathway analysis of these differentially regulated gene sets showed strong dampening of interferon response pathways by RTX treatment in both cell types (**Fig. S13A**, **Table S5**).

**Figure 6.**
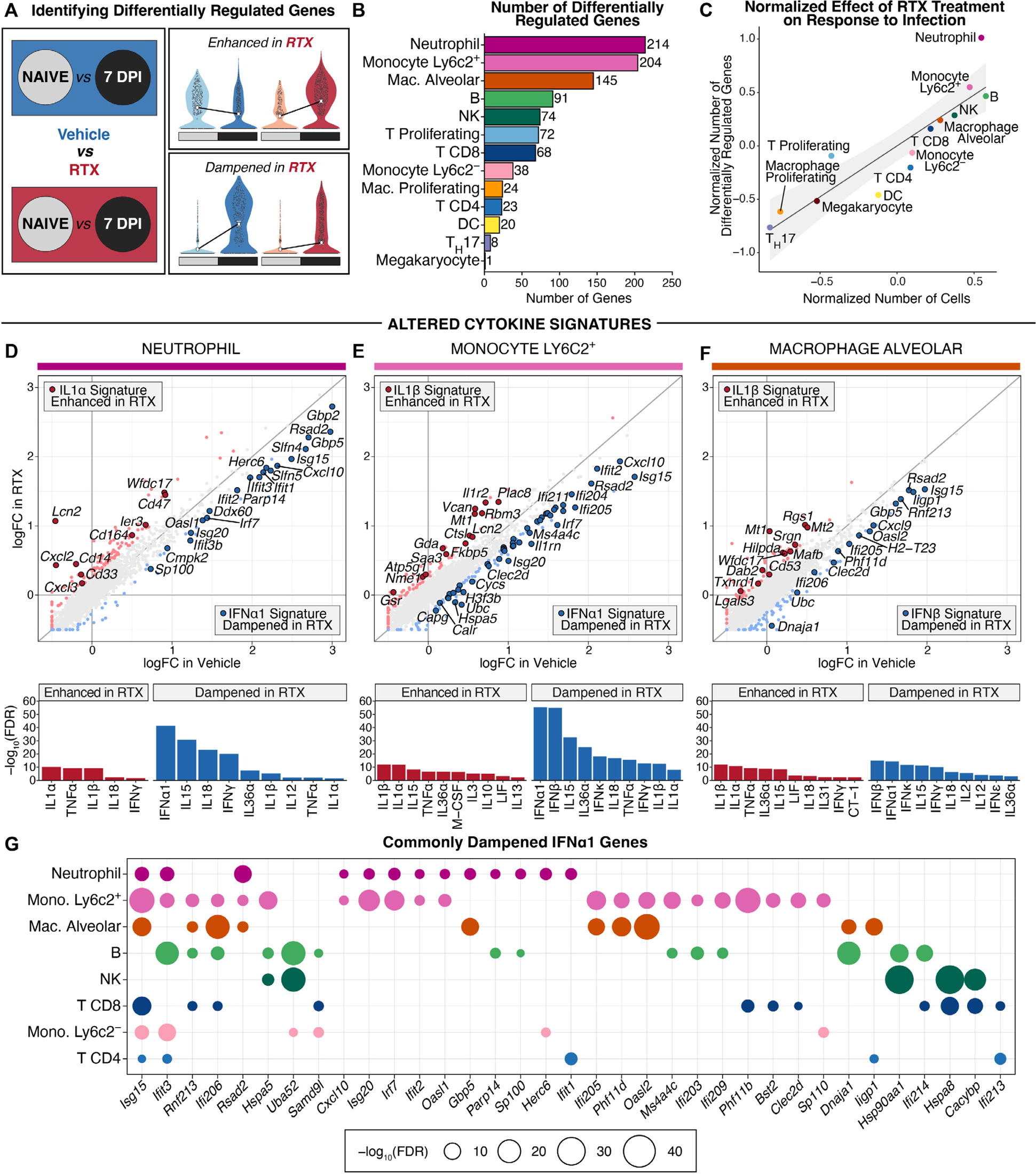
Differentially regulated genes indicate RTX treatment-induced shifts in cell-type specific cytokine responses. **(A)** Schematic representation of the comparisons (left) used to identify viral response genes modified by RTX treatment; example violin plots (right) to illustrate potential ways a gene can be differentially regulated in RTX compared to vehicle. **(B)** Bar plot indicating the number of differentially regulated genes (x axis, label) identified for each cell type (y axis) with at least one differentially regulated gene. **(C)** Scatter plot showing the relationship between the number of cells (x axis) and the number of differentially regulated genes (y axis) (log_50_ scaled) after regressing out the effects of differences in the average number of genes expressed by each cell type (labels). Axes show the residual values from the line of best fit between the log average number of genes expressed and the log number of cells (x axis) or log number of differentially regulated genes (y axis) **(D-F).** Scatter plot (top) of all genes showing the log_50_ fold change with infection (7 DPI vs Naïve) in Vehicle (x axis) compared to the log_50_ fold change with infection in RTX (y axis). Points for differentially regulated genes are colored lighter blue if significantly (adjusted *limma* p value < 0.05) dampened in RTX compared to vehicle and lighter red if significantly enhanced in RTX; gray points are genes that change a similar amount with infection in vehicle and RTX. Larger points with black outline are differentially regulated genes corresponding to the top cytokine result dampened (dark blue) or enhanced (dark red) in RTX. Bar plot (bottom) indicates the −log_50_ adjusted hypergeometric p value (y axis) for the overlap of the genes enhanced (left, red bars) or dampened (right, blue bars) in RTX with cell-type specific gene signatures of individual cytokines (x axis). Gene scatter plot and cytokine significance bar plots shown for **(D)** neutrophils, **(E)** *Ly6c2*+ monocytes, and **(F)** alveolar macrophages. **(G)** Dot plot showing IFN*α*1-induced genes (x axis) that were also significantly enriched among the genes dampened by RTX treatment in two or more cell types (y axis, color). Dots are only shown when a gene is significantly (adjusted *limma* p value < 0.05) dampened in RTX compared to vehicle and is a IFN*α*1-induced gene for that cell type; dot size indicates the −log_50_ adjusted *limma* p value of that gene for that cell type (legend).

Given the critical role of cytokines in immunity to viral infections, we tested whether these differentially regulated genes could be used to identify the upstream cytokines driving perturbed responses after nociceptor ablation. We tested for overlap with a recently published database of cell type-specific transcriptional signatures induced by in-vivo treatment with individual cytokines (47) (**Table S5**). This approach identified significant enrichment of cell-type specific responses to type I interferons, particularly IFNα1 and IFNβ, and the interferon-associated cytokine, IL-15, as top hits among the RTX-dampened genes across almost all cell types, whereas IL-1α, IL-1β, and TNF*α*-induced signatures consistently overlapped with the RTX-enhanced genes of different cell types (**Fig. 6D-F** and **Fig. S13B**). Among the IFNα1-induced genes previously identified (47), 35 genes contributed to significant enrichment of this cytokine’s signature across multiple cell types (**Fig. 6G**). The ubiquitin-like modifier *Isg15*, inhibiter of cellular and viral processes *Ifit3*, atypical E3 ubiquitin ligase *Rnf213*, and transcriptional regulator *Ifi206* were the most broadly shared genes, and the strongest induction of this signature was observed in neutrophils and *Ly6c2*^+^ monocytes. Overall, this suggests that nociceptor ablation interferes with the normal induction of interferon signaling in immune cells upon infection with IAV.

### Classification of pulmonary neutrophil subsets in the setting of influenza infection

Immune profiling by flow cytometry and scRNAseq demonstrated that neutrophils in nociceptor-deficient mice were both significantly expanded (**Fig. 4B** and **Fig. S7**) and transcriptionally perturbed during influenza infection (**Fig. 6**). To better understand these changes, we isolated and re-clustered all 2,921 neutrophils, revealing three distinct states (**Fig. 7A**, **Table S6**). *Thbs1*, a gene encoding the adhesive glycoprotein thrombospondin 1, was the top marker of the cluster of 1,093 naïve lung neutrophils (**Fig. 7A-B** and **Fig. S14A**). The neutrophils in the lung at 7 DPI formed two clusters: a smaller population of 335 neutrophils that specifically expressed *Stfa2l1*, a member of the cystatin family of cysteine protease inhibitors, and a larger cluster of 1,493 neutrophils expressing *Nmes1* (also known as *C15orf48* and *Aa467197*) which encodes a mitochondrial protein linked to autophagy and inflammasome activation (48, 49) (**Fig. 7A-B** and **Fig. S14A**). Less than half (41.7%) of the cells in the cluster of naïve *Thbs1*^+^ neutrophils were from RTX-treated mice and of the two neutrophil populations present in the lungs at 7 DPI, cells from RTX were a minority (35.8%) of the *Stfa2l1*^+^ population, whereas the *Nmes1*^+^ neutrophils were predominately (67%) from nociceptor-deficient mice (**Fig. 7C** and **Fig. S14B**). This suggests that the increase in neutrophils in RTX-treated mice post-IAV infection more specifically reflects an expansion of the *Nmes1*^+^ neutrophil population.

**Figure 7.**
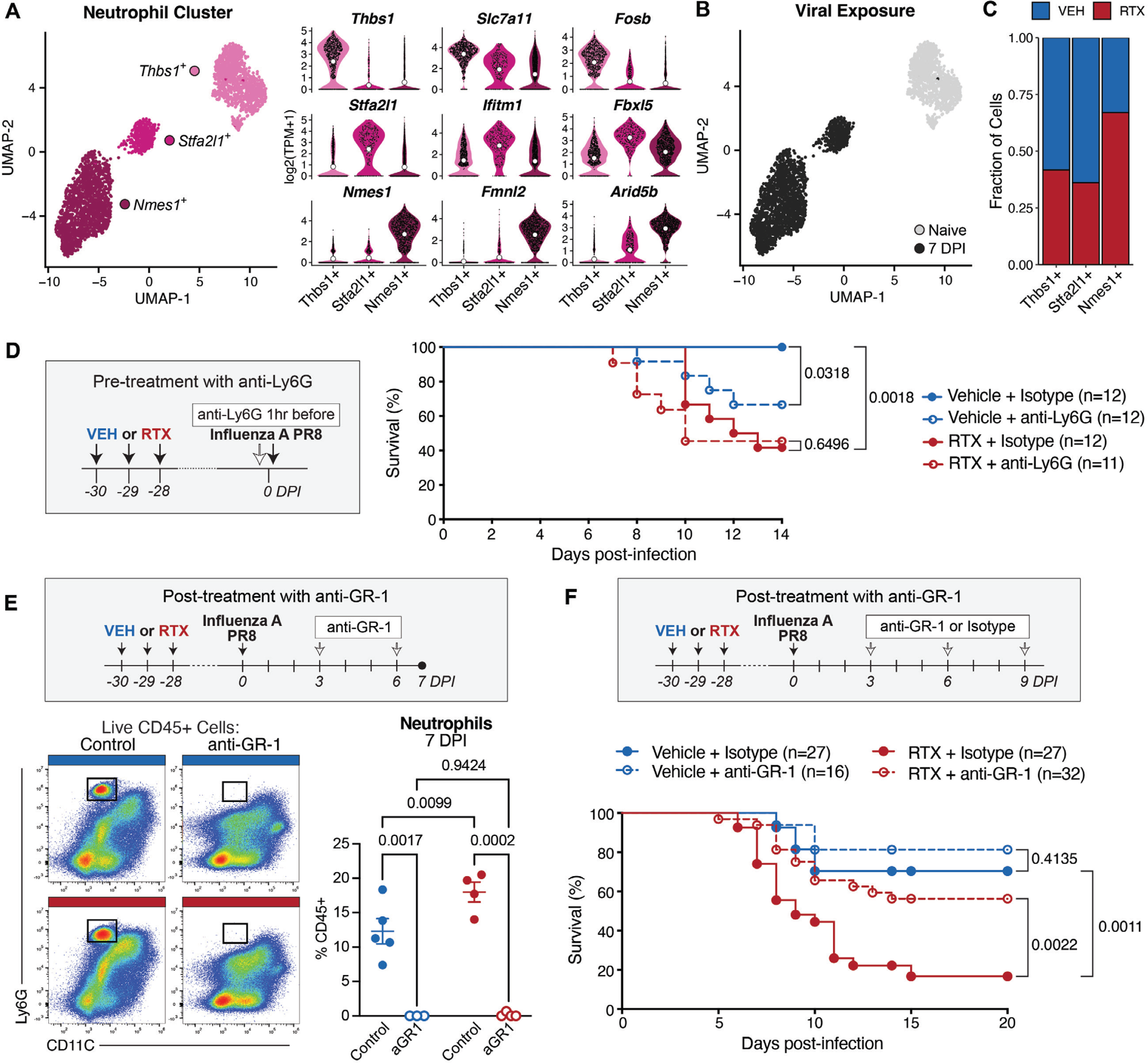
Anti-GR-1 treatment rescues nociceptor-ablated mice from influenza infection-induced mortality. **(A)** UMAP (left) of 2,921 neutrophils colored and labeled by the top marker of each cluster. Violin plots (right) show expression (y axis) of the top 3 markers specific to each neutrophil cluster (x axis, color). **(B)** UMAP of neutrophils colored by viral exposure group. **(C)** Cumulative bar plot showing the fraction of cells (y axis) in each neutrophil cluster (x axis) from vehicle or RTX treatment groups (color legend). **(D)** Mice were treated with vehicle or RTX and then 4 weeks later pre-treated with intraperitoneal injection of anti-Ly6G (50µg) or isotype control an hour before intranasal inoculation with IAV PR8 (schematic). Survival was monitored after infection and is shown for the four treatment groups (legend). (n=11-12 per group). **(E)** Vehicle and RTX treated mice were infected intranasally with IAV PR8 then post-treated with intraperitoneal injections of anti-GR-1 (250µg per mouse) at days 3 and 6 post-infection and lungs collected for flow cytometry at 7 DPI (schematic). Flow cytometry plot (left) of live CD45^+^ lung immune cells from control and anti-GR-1 injected vehicle and RTX treated mice (color bar) gated for neutrophils (Ly6G^+^CD11C^−^), quantified as a percentage of CD45+ cells (right). (n=4-5 per group). **(F)** Vehicle and RTX treated mice were infected intranasally with IAV PR8 then received intraperitoneally injected with 250µg per mouse of anti-GR-1 or isotype control at days 3, 6, and 9 post-infection (schematic) then monitored for survival. Data shown is pooled from three independent experiments conducted with mice of both sexes. (n=16-32 mice per group). Log-rank test was used in (D and F); Student’s t test in (E); error bars show mean±SEM; P values labeled.

We then used principal component analysis (PCA) and correlations to better understand differences between the three neutrophil states (**Fig. S14C-D**). The first principal component distinguished *Nmes1*^+^ neutrophils from the other two clusters and the second separated the *Stfa2l1*^+^ and *Thbs1*^+^ states (**Fig. S14C**), indicating the *Nmes1*^+^ was most distinct. Consistently, correlation analysis showed that the *Thbs1*^+^ and *Stfa2l1*^+^ neutrophils were relatively similar to one another, whereas the *Nmes1*^+^ neutrophils were negatively correlated with the other two clusters, including a strong negative correlation with the *Thbs1*^+^ cluster of naïve neutrophils (**Fig. S14D**).

We next tested whether the cytokine signatures altered by RTX treatment were perturbed within specific neutrophil subsets. Of the IFN *α* 1-induced genes found to be dampened in neutrophils, we see little expression in the *Thbs1*^+^ cluster of naïve neutrophils and upregulation in both the *Stfa2l1*^+^ and *Nmes1*^+^ clusters from 7 DPI, illustrating the induction by influenza infection in both RTX and vehicle-treated mice (**Fig. S14E**). Interestingly, of the IFN*α*1 signature genes, 4 are significantly dampened in RTX *Stfa2l1*^+^ neutrophils, while 9 genes showed a reduction in RTX *Nmes1*^+^ cells, suggesting that the deficit in interferon signaling observed overall for neutrophils is present across subsets and most striking within the *Nmes1*^+^ neutrophils (**Fig. S14E**). We then looked for expression of specific genes encoding cytokines that were altered by RTX treatment and found that both *Il1a* and *Tnf,* cytokines whose signatures were enhanced in RTX, were highly and specifically expressed by the *Nmes1*^+^ cluster of neutrophils (**Fig. S14F**), linking the perturbed cytokine signaling and switch from interferon to IL-1a/IL-1b/TNF*α* to an expansion of the *Nmes1*^+^ neutrophil state.

### Myeloid cell depletion rescues nociceptor-deficient mice from influenza-induced mortality

Given the compositional and transcriptional changes we observed in the lung immune compartment of nociceptor deficient mice, we hypothesized that immune dysregulation in RTX-treated mice was driving increased mortality after influenza infection. We first tested whether neutrophils, specifically, played a role in survival from influenza infection by pre-treating RTX- and vehicle-treated mice with 50μg of anti-Ly6G antibody or isotype control one hour before intranasal inoculation with IAV PR8. Depleting neutrophils prior to infection via pre-treatment with anti-Ly6G had no effect on the survival of RTX-treated mice, whereas anti-Ly6G pre-treated vehicle mice exhibited worsened survival compared to the isotype control (**Fig. 7D**). This data likely reflects the importance of neutrophils in the initial host immune response to influenza (50).

Our flow cytometry and scRNAseq datasets revealed a broader involvement of multiple myeloid populations, with significantly increased neutrophils and monocytes in BALF (**Fig. S7**), and expansion of both neutrophils and monocyte-derived macrophages in lung tissues (**Fig. 4** and **Fig. 5**). In addition to neutrophils, *Ly6c2*^+^ monocytes also exhibited substantial transcriptional changes and may play a major role in immunopathology (**Fig. 6**). Thus, we chose to target myeloid cells more broadly by treating mice with 250µg of anti-GR-1 antibody, which depletes both circulating Ly6G^+^ neutrophils and Ly6C^+^ monocytes (51). Of note, a recently described population of influenza-induced Ly6G^+^ monocyte-derived macrophages (44) may also be targeted by anti-GR-1 treatment. To preserve the beneficial early roles of neutrophils, we administered the antibody after infection.

We performed intraperitoneal injections of anti-GR-1 antibody or isotype control antibodies at 3, 6, and 9 DPI in both RTX-treated and vehicle-treated mice (**Fig. 7E-F**). By flow cytometry, we confirmed that following the first two anti-GR-1 injections, neutrophils were significantly depleted in both vehicle and RTX-treated mice at 7 DPI (**Fig. 7E**). Anti-GR-1 also reduced the proportion of monocytes, while CD11B^+^F4/80^+^Ly6C^Lo^CD11c^Hi^ cells were increased as a fraction of all immune cells (**Fig. S15**). Following infection, we found that anti-GR-1 treatment significantly improved survival in RTX-treated mice compared to isotype-treated RTX mice, whereas there was no significant difference between anti-GR-1 and isotype-treated vehicle mice (**Fig. 7F**). These results indicate that RTX-induced immunopathology drives mortality in influenza infection and that targeting myeloid cells can mitigate this effect in nociceptor-deficient mice.

## DISCUSSION

Host defense against respiratory viral pathogens requires a delicate balance between defensive efforts to control the spread of the virus and preserving tissue integrity and lung function (4). Our study identifies a key role for TRPV1^+^ sensory neurons in striking this balance; we demonstrate that nociceptors are critical to limit pathological inflammation, lung damage, and mortality from influenza A pneumonia. In the absence of TRPV1^+^ sensory neurons, influenza infection results in dysregulated immune responses, including altered proinflammatory cytokine induction and increased neutrophil and monocyte-derived macrophage numbers, which significantly worsened survival without affecting viral transcript levels or body weight loss. The protective role of nociceptors was not mediated by local release of neuropeptides CGRP or substance P within the lungs. Instead, nociceptor-deficient mice were rescued by depleting myeloid cells after infection, underscoring the impact of poorly controlled inflammation on survival from influenza infection.

This study builds upon existing literature highlighting the diverse roles of airway-innervating sensory neurons in the response to inhaled pathogens. A recent study employed vagotomy, pharmacological inhibition with the sodium channel blocker QX-314, and genetic depletion of neurons with *Phox2b*-Cre mice to explore the role of vagal sensory neurons in influenza infection (28). Consistent with our findings, the authors reported that ablation of vagal sensory neurons worsened clinical features and survival from influenza infection without affecting viral burden (28). However, there are differences with our current study. The aforementioned study targeted the majority of nodose neurons (*Phox2b*^+^) or used tools that silenced both vagal efferent and afferent neurons (vagotomy, QX-314). In contrast, our study employed more specific genetic and pharmacological approaches to target only nociceptors (*Trpv1*^+^ and/or Nav1.8^+^). We also observed significant dysregulation of immune responses in the lungs of nociceptor-ablated mice and no weight loss during infection, which differs from the previous study.

Of potential relevance, *Gabra1*^+^ petrosal neurons were recently identified as airway-innervating neurons that modulate appetite, weight loss, and sickness behaviors following PR8 infection (10). *Trpv1*^+^ neurons do not express *Gabra1* (**Fig. S3**). This distinction may explain why we did not observe effects on food or water consumption or body weight loss. Our study identified a robust effect of nociceptor ablation on the immune compartment of the lung during infection, particularly with an increase in myeloid cells such as neutrophils and monocyte-derived macrophages. The fact that broader ablation of all vagal sensory neurons did not result in similar immune changes (28) suggests that nociceptors work in tandem with other, currently unappreciated populations of vagal sensory neurons to achieve an appropriate balance in the immune response to IAV infection.

Contrary to the protective role of nociceptors in IAV infection, previous work from our group showed that nociceptor ablation improved survival from lethal *S. aureus* infection (11). In the context of bacterial pneumonia, nociceptor-deficient mice exhibited an increase in neutrophils and γδ T cells, coinciding with a reduction in bacterial burden and a survival benefit for the host, which was dependent on CGRP signaling (11). The dichotomy between the effects of nociceptor ablation on host response to bacterial lung infection versus influenza virus infection underscores the pleiotropic roles of both nociceptors and neutrophils, which must be recognized for their beneficial and detrimental effects in a context-dependent manner.

We present the first single-cell transcriptomic dataset examining the impact of neuronal ablation on the lung immune compartment at homeostasis and post-infection. Our scRNAseq data reveal a reduction in interferon signaling and an increase in IL-1 and TNFα signaling in RTX-treated mice. This shift mirrors the known role of the type I interferon-IL-1 axis in driving immunopathology (52–54). Typically, a deficit in IL-1 signaling coupled with increased interferon responses worsens bacterial infections, while the opposite is true for viral infections (52). This imbalance might explain why nociceptor-deficient mice are more resistant to bacterial infections like *S. aureus* but have worse outcomes with influenza. Supporting these ideas, we observed delayed and lower IFNα levels in BALF of RTX-treated mice post-IAV infection, along with elevated G-CSF levels, which may contribute to the increased neutrophils and monocyte-derived macrophages at 7 DPI. Closer inspection of the neutrophils in our scRNAseq dataset revealed three distinct populations: lung-resident neutrophils, which have been described previously (55–57), and two infection-induced states. Interestingly, the top marker for naïve lung neutrophils was *Thbs1*, a gene encoding thrombospondin-1, which was recently shown to be induced by immune cells, promoting tissue repair in the skin (58) and limiting pain via interaction with skin-innervating nociceptors (59). It remains to be seen whether *Thbs1* in neutrophils may play similar protective roles in the lungs at baseline.

Neutrophil depletion via anti-Ly6G treatment before infection had no impact on the survival of RTX-treated mice, but worsened survival in vehicle-treated controls, suggesting that lung-resident neutrophils are protective early in IAV infection. The two clusters of neutrophils present in the lungs after IAV infection, were specifically marked by the genes *Stfa2l1* and *Nmes1*, with RTX treatment pushing the balance from the *Stfa2l1*^+^ subset to the *Nmes1*^+^ state. Based on their transcriptional similarities, we suspect that the *Stfa2l1*^+^ cluster of neutrophils may reflect a similar population to the *Thbs1*^+^ neutrophils present in the lungs at baseline which have undergone significant changes in response to IAV infection, while the *Nmes1*^+^ neutrophils are likely only recruited to the lung after infection. The *Nmes1*^+^ neutrophils are high expressors of *Il1a* and *Tnf*, situating them as a source of the pro-inflammatory cytokines in RTX-treated mice. We demonstrate that myeloid cell depletion with anti-GR-1 treatment at days 3, 6, and 9 post-infection efficiently depleted neutrophil and monocyte-derived populations within the lung and significantly improved survival in nociceptor-deficient mice, illustrating the injurious role that myeloid cells play in driving increased mortality from influenza infection in nociceptor-deficient mice.

In summary, we demonstrate a previously unrecognized role of vagal TRPV1^+^ sensory neurons in modulating host defense to influenza infection. Nevertheless, some outstanding questions should be addressed by future research. While we ruled out a role for CGRP and substance P, it is not clear whether local release of other neuronal mediators could regulate the myeloid response and protect against pathology. It is also unclear whether there are vagal-brain-autonomic feedback circuits that could be suppressing the immunopathology. Recent work has shown that in mouse models of LPS-induced septic shock, specific cytokines induce activation of distinct vagal and brainstem neurons; in turn, these neurons signal back to the body via efferent circuits to regulate inflammation in the peripheral tissues (13). Similar airway-to-brain circuits have also been defined in the contexts of *P. aeruginosa* infection (26) and allergic airway inflammation (60). It is possible that a related or distinct circuit is activated by influenza infection through TRPV1^+^ neurons. While we demonstrated that nociceptors are critical regulators of myeloid cell recruitment to the lung during influenza A infection, it remains unknown whether specific cell types signal to TRPV1^+^ sensory neurons to drive this phenotype. How neurons interact with lung-resident cells to maintain airway homeostasis and during inflammation is also unknown. Finally, the roles of downstream neurons activated by vagal nociceptors and the contributions of a lung-brain axis via vagal efferent signaling has yet to be determined in the case of viral infection.

## MATERIALS AND METHODS

### Mice

C57BL/6, B6.129-*Trpv1*tm1(cre)Bbm/J, B6.*Rosa26*-stop(flox)-DTA, B6.129X1-*Trpv1*tm1Jul/J (Trpv1 KO), B6.Cg-*Tac1*tm1Bbm/J were purchased from Jackson Laboratories. Nav1.8-Cre mice were provided by J. Wood (University College London). *Calca*^−/−^ mice (61) were provided by V. Kuchroo (Harvard Medical School). *Trpv1-*DTR mice were a gift from Mark Hoon (NIH). For Nav1.8-lineage neuron depletion experiments, Nav1.8-Cre^+/−^ mice were crossed with *Rosa26*-stop(flox)-DTA^+/+^ mice to generate Nav1.8-lineage neuron-depleted Nav1.8-Cre^+/−^;DTA^+/−^ (Nav1.8^DTA^) mice and control Nav1.8-Cre^−/−^;DTA^+/−^ (Control) littermates. For Calca and Substance P deficient mouse experiments, the knockout mice were cohoused with age- and sex-matched B6 mice for at least three weeks before experiments. Age-matched 6- to 12-week-old littermate male and female mice were used for experiments.

At Harvard Medical School (HMS), mice were bred and housed in a full-barrier, specific pathogen free animal facility. All animal experiments performed at Harvard Medical School were approved by the HMS Institutional Animal Use and Care Committee and Committee on Microbiological Safety. At Yale University, mice were maintained in a specific pathogen-free facility and all animal experiments were performed in accordance with institutional regulations after protocol review and approval by Yale University’s Institutional Animal Care and Use Committee. At New York University Langone Health Center, all mice were bred in-house and maintained with food and water ad libitum under a 12-hour dark/light cycle in a pathogen-free facility. All experiments were performed with approval by the New York University Langone Health Center Institutional Animal Care and Use Committee and in accordance with guidelines from the National Institutes of Health, the Animal Welfare Act, and the U.S. Federal Law. All influenza experiments were performed with biosafety level 2 precautions and within dedicated areas of animal facilities.

### Influenza A virus infections

For experiments at Harvard Medical School and at NYU, Influenza A virus Puerto Rico strain 8 (PR8) was purchased from Charles River Laboratories. WT, RTX-treated, Nav1.8^DTA^, and Trpv1-DTR mice were inoculated with 50 TCID_50_(see below) PR8. For Calca^−/−^ and Tac1^−/−^ experiments, mice were infected with 50 TCID_50_ PR8, as well as higher doses of 250 and 500 TCID_50_, respectively. For experiments at Yale University, IAV WSN was similarly diluted in saline, and mice infected intranasally with 300 plaque forming units (PFU) or 250 PFU per mouse. For experiments at NYU, 50 Egg-infectious dose (EID_50_) or 75 EID_50_ PR8 IAV inoculated into mice, with half a dose administered per nostril.

### TCID_50_ Hemagglutination Assay

MCDK cells (ATCC) were cultured in DMEM supplemented with 10% FBS, 1X Penicillin/Streptomycin, and 2mM L-glutamine. MDCK cells were seeded in 96-well dishes overnight. The next day, cells were washed twice with PBS and infected with half-log dilutions of virus in 200μl inoculum. Viral infections were performed in UltraMDCK media (Lonza) supplemented with 1X Penicillin/Streptomycin and TPCK trypsin at 1ug/mL (Sigma). Infected cells were incubated at 37°C for 48 hours. Cell culture supernatant was then added to V-bottom plates on ice. 0.5% chicken red blood cells (Rockland) in PBS were added to the cell culture supernatant 1:1 and incubated on ice for 1 hour. Wells were then assessed for agglutination and TCID_50_ was calculated utilizing the Reed–Muench Method (62).

### Influenza A Virus WSN Plaque Assay

Infectious virions of WSN (A/WSN/1933) were quantified by plaque assay in MDCK cells (culture conditions described above). Cells were seeded to confluence in 6-well plates. MDCK cells were infected with log-fold viral dilutions in 0.1% BSA in PBS at 37°C for one hour. Cells were washed with PBS and overlayed in 1X MEM (Gibco) with 1ug/mL TPCK trypsin (Sigma), 0.5% NaHCO_50_, 1X Penicillin/Streptomycin, 1% agarose, 2.1% BSA (Fisher). Inverted plates were cultured at 37°C for 72 hours and then fixed with 4% PFA, subsequently stained with 0.1% crystal violet in 20% ethanol, and plaques were counted.

### Viral NP transcript measurement

Transcriptional viral load was determined by measuring the influenza virus nucleoprotein (NP) in BALF samples as previously reported (10). Briefly, the mouse neck was surgically opened and 1ml of PBS was inserted into the trachea and aspirated back (three times, in total 3 ml was collected). BALF cells were isolated by centrifuge at 300g for 5 min, and resuspended in 1 ml Trizol (15596026, ThermoFisher) for RNA extraction and following cDNA synthesis according to the manufacturer’s instructions. Influenza viral nucleoprotein primers (forward GACGATGCAACGGCTGGTCTG and reverse ACCATTGTTCCAACTCCTTT) were used for qPCR and GAPDH (forward ACAGTCCATGCCATCACTGCC and reverse GCCTGCTTCACCACCTTCTTG) was used for the normalization.

### Nociceptor neuron depletion

Systemic nociceptor neuron ablation by RTX treatment was performed as previously described (11, 31, 32). Briefly, 4-week-old C57BL/6 mice were anesthetized with isoflurane and injected subcutaneously in the flank with escalating doses (30, 70, 100 mg/kg on consecutive days) of resiniferatoxin (RTX, Alomone Lab) or vehicle (2% DMSO/0.15% Tween-80/PBS). Mice were allowed to rest for 4 weeks before experiments. Loss of nociceptor neurons was confirmed by reduced thermal responses to noxious heat during hot plate tests. Systemic nociceptor neuron ablation of Trpv1-Dtr mice by diphtheria toxin (DT) treatment was performed as described (11, 37). Briefly, the Trpv1-Dtr mice were injected intraperitoneally with 200 ng of DT (Sigma Aldrich) dissolved in 100 μL PBS or with 100 μL PBS (vehicle) daily for 21 days. For vagal ganglia-targeted ablation, bilateral intraganglionic injections of DT or PBS into Trpv1-Dtr mice was performed as described (11, 22). 20 ng DT in 120 nL PBS containing 0.05% Fast Green was injected into nodose/jugular/petrosal VG with a nanoinjector (Drummond Scientific Company).

### AAV injections for labeling of TRPV1^+^ lung-innervating neurons

For visualization of lung innervation, Trpv1-Cre mice were intraperitoneally injected on postnatal day 1 with 10uL of 2×10^11^ genome copies/mL AAV9-FLEX-tdTomato (Addgene, 8306-AAV9) then sacrificed for lung harvest at 6-8 weeks old. For anterograde labeling of vagal TRPV1^+^ neurons that innervate the lungs, we performed bilateral intraganglionic injections of AAV9-FLEX-tdTomato into the vagal nodose/jugular ganglia of mice (Penn Vector Core, AV-9-ALL864) into Trpv1-Cre mice as previously described (11, 30). Mice were anesthetized with 1–3% isoflurane with oxygen. The nodose/jugular/petrosal ganglia were exposed and injected sequentially after a midline incision in the neck (*∼*1.5 cm in length). A Nanoject II (Drummond Scientific) was used to inject 120 nL of viral solution (1.3×10^13^ genome copies/mL in PBS with 0.05% Fast Green). Two weeks after AAV injection, mice were sacrificed and whole lungs were harvested for sectioning and immunostaining.

### Immunostaining

Sacrificed mice were intracardially perfused with cold PBS, then lungs were collected and fixed in 4% PFA/PBS at 4°C overnight. 200μm thick precision-cut lung sections were obtained using a compresstome® (Precisionary Instruments), collected into blocking solution (PBS + 0.5% Triton X-100 + 5% Donkey Serum) and allowed to incubate overnight shaking at 4°C. Sections were then incubated with goat anti-tdTomato (Origene, AB0040-200, 1:500 dilution), mouse anti-βIII-tubulin (Abcam, ab7751, 1:500 dilution), and rat anti-E-cadherin (Invitrogen, 13-1900, 1:500 dilution) diluted in blocking solution, shaking at 4°C for three nights. Sections were washed with blocking solution, then transferred to secondary antibodies Alexa 594 donkey anti-goat (Abcam), Alexa 488 donkey anti-mouse (Abcam), and Alexa 647 donkey anti-rat (Abcam) at a 1:250 dilution in blocking solution and allowed to incubate overnight shaking at 4°C. Sections were then counterstained with DAPI in blocking solution, and post-fixed in 4% PFA/PBS for 1 hour, then washed with PBS. Images were obtained using a Stellaris 8 FALCON CFS system (Leica).

For ganglia immunostaining, mice were perfused with 4% paraformaldehyde (PFA) in PBS after euthanasia. Vagal ganglia were dissected, fixed in 4% PFA/PBS at 4 °C for 1 hour and incubated with 30% sucrose/PBS at 4 °C overnight before embedding in optimal cutting temperature compound (OCT, Tissue-Tek, PA). 20 μm cryosections were cut and immunostained with the following antibodies: guinea pig anti-TRPV1 (Millipore, AB5566, dilution 1:500), rabbit anti-ATF3 (Sigma, HPA001562, dilution 1:500), and mouse anti-βIII-tubulin (Abcam, ab7751, dilution 1:500). Secondary antibodies included CF-488A goat anti–guinea pig IgG (Sigma, 1:500), Alexa 647 donkey anti-mouse IgG (Abcam, 1:500), and Alexa 594 donkey anti-rabbit IgG (Abcam, 1:500). Sections were mounted in Vectashield and imaged with a Stellaris 8 FALCON CFS system (Leica).

### Vagal ganglia histological analysis

Following staining for TRPV1, ATF3, and Tuj1 and imaging described above, the number of ATF3^+^ cells, TRPV1^+^ cells and Tuj1^+^ cell was quantified in Fiji ImageJ using a customized Macro applied to all images. The fraction of ATF3^+^ neurons was calculated as the number of ATF3^+^ cells divided by the number of Tuj1^+^ cells and the ATF3^+^TRPV1^+^ fraction as the number of ATF3^+^TRPV1^+^ cells divided by the total number of TRPV1^+^ cells.

### H&E staining

Lung tissue was collected directly after sacrifice or following intracardial perfusion with cold PBS, as indicated, then washed with PBS and fixed with 4% PFA/PBS overnight. Paraffin-embedded sections of the lungs were subjected to H&E staining and then examined by Thunder 3D Cell Culture (Leica).

### Pathology Scoring

H&E images of non-perfused lung tissue sections were scored by a blinded reviewer, following previously described lung histology scoring criteria (63). Briefly, for each image 5 fields of view from throughout the tissue were selected and evaluated for the degree of edema, hemorrhage, cell infiltration, and airway necrosis; the score presented for an image represents the average of the scores for the different fields of view. The minimum score of 0 represents normal healthy lung tissue and a maximum score of 4 corresponds to increased pathological features throughout the lung.

### Vital sign measurements

Vital signs (oxygen saturation, heart rate, and breathing rate) were measured in conscious mice using collar-based mouseOx pulse oximeter (Starr Life Sciences). Briefly, fur around the collar area of mice was first removed prior to vital sign measurements using Nair hair remover. To insert collar sensor clamp into the neck area of mouse at the time of measurement, mice were lightly anesthetized with 2-3% isoflurane. Then, vital-sign measurements were recorded after mouse became fully awake and mobile using MouseOX software (Starr Life Sciences). For accuracy, we measured the vital signs every 10 seconds for a total time of 5 minutes, and then reported the average of all measurements.

### Bronchoalveolar lavage fluid collection and processing

Collection and analysis of bronchoalveolar lavage fluid (BALF) was conducted as described previously (11). Briefly, the trachea was exposed and cannulated with a 20-gauge catheter. BALF was collected by instilling 0.8mL of cold PBS containing heparin and dextrose, then centrifuged for 7 minutes at 4,000 RPM and 4°C to separate the supernatant and cell pellet. For protein and cytokine analysis, the cell-free BALF supernatant was filtered through a 0.22um filter and mixed with protease/phosphatase inhibitor cocktail. For isolation of immune cells from BALF, the cell pellet was briefly incubated in red blood cell lysis buffer, then processed for flow cytometry.

### Cytokine measurements

BALF supernatants were collected from the RTX- and Vehicle-treated mice at the indicated time points post IAV infection. The supernatant samples were used for cytokine/chemokine detection by Mouse Cytokine/Chemokine 44-Plex Discovery Assay® Array (MD44, Eve technologies) targeting 44 cytokines and chemokines. Levels of IFN*α* and IFN*λ* were measured using enzyme-linked immunosorbent assay (ELISA) kits according to the manufacturer’s instructions (R&D Systems).

### Lung immune cell isolation

Mice were sacrificed and intracardially perfused with PBS before lung collection. Lung tissues were minced with scissors then digested with 1 mg/ml Collagenase A (10103586001, Sigma) and 100 U/ml DNase I (Sigma) in a shaker at 37°C 250 rpm for 1 hour to obtain single-cell suspensions. The cell suspensions were first depleted of red blood cells and then separated into two parts via 40%/80% Percoll gradient centrifugation at 1200g for 20 min. After centrifugation, cells between the two layers were identified as immune cells and used for flow staining.

### Flow cytometry staining

Immune cells isolated above were stained with the following antibodies (Live/Dead Fixable NIR, Invitrogen, #L34975, 1:500; CD45, Pacific Blue, Biolegend, #103126, 1:500; SiglecF, PE, BD, #552126, 1:300; CD11C, FITC, BioLegend, #117306, 1:300; F4/80, Percp cy5.5, BioLegend, #123128, 1:300; CD11B, BV605, Biolegend, #101237, 1:300; Ly6G, PEcy7, Biolegend, #127617, 1:300; Ly6C, APC A700, BioLegend, #128024, 1:300; CD19,BV650, BioLegend, #115541, 1:300; CD3, BV605, BioLegend, #100237, 1:300; CD4, AF700, BioLegend, #100536, 1:300; CD8, Percp cy5.5, Biolegend, #100733, 1:300; GATA3, PEcy7, eBioscience, #25-9966-42, 1:300; RoRgt, APC, eBioscience, #17-6981-80, 1:300; Foxp3, FITC, Biolegend, #126405, 1:300; NK1.1, PE, Biolegend, #108707, 1:300) on ice for 20 minutes in flow buffer (2% FBS, 2mM EDTA in PBS) and washed twice with flow buffer. Cells were permeabilized and stained using a Fixation/Permeabilization kit (BD) according to the manufacturer’s instructions for intranuclear staining. Data were acquired on a CytoflexS flow cytometer (Beckman Coulter) and analyzed in Flowjo v10.

### Flow cytometry of CD4 and CD8 T cells and NP tetramer-specific T cells

Immune cells were isolated as described above then stained with the following antibodies (Live Dead AF350 NHS Ester, Thermofisher, #A10168, 1:400; CD62L-BUV737, BD Bioscience, #612833, 1:100; NK1.1-BV711, Biolegend, #108745, 1:100; CD3-BV421, Biolegend, #100228, 1:100; CD45-BV510, Biolegend, #103138, 1:100; CD4-BV650, Biolegend, #100546, 1:200; CD8a-BV785, Biolegend, #100750, 1:200; KLRG1-FITC, ThermoFisher, #11-5893-82, 1:100; CD103-PerCP-Cy5.5, Biolegend, #121416, 1:100; TCRgd-PE, Biolegend, #118108, 1:100; CD127-PE-Dazzle594, Biolegend, #135032, 1:100; CD11a-PE-Cy7, Biolegend, #101122, 1:100; CD69-AF700, Biolegend, #104539, 1:100; CD44-APC-Cy7, Biolegend, #103028, 1:200; and NP Tetramer-APC, NIH Tetramer Core, 1:200) for 20 min in 4°C. Cells were fixed with 2% PFA for 20 min in 4°C and resuspended in FACS buffer. Cells suspension processed on Bio-Rad ZE5 Yeti instrument and data analyzed in FlowJo software.

### Isolation of CD45^+^ immune cells for single-cell RNA sequencing

Mice were sacrificed by carbon dioxide followed by cervical dislocation and were immediately perfused transcardially with 10ml PBS solution. Collected lung lobes were rinsed in 10% FBS/RPMI and minced into small pieces. Samples were digested (100µg/ml DNase and 0.4 mg/ml collagenase in 10% FBS/RPMI) using GentleMacs Dissociator (Miltenyi biotec), followed by filtering with 70μm strainer using 10% FBS/RPMI. Samples were then labeled with CD45 MicroBeads (Miltenyi Biotec) at 4°C for 15 minutes, washed with MACS buffer (Miltenyi biotec), and filtered with 40μm strainer. The resulting cell suspensions were then used for magnetic-activated cell sorting (MACS)-based purification using MS MACS columns (Miltenyi Biotec) following the manufacturer’s directions. Cells were resuspended in 0.04% BSA/PBS and pooled from two mice for each of the naïve samples (vehicle and RTX), and from three mice for each of the 7 DPI samples (vehicle and RTX) to generate a total of four cDNA libraries, which were sequenced on an Illumina NovaSeq with Nextera XT assay.

### Myeloid cell depletion

For the depletion of neutrophils in vehicle- and RTX-treated mice, we injected mice with anti-Ly6G antibody at 1 hour prior to PR8 infection. Briefly, mice were injected intraperitoneally with a single dose of 50 µg of anti-mouse Ly6G monoclonal antibody (clone 1A8, Bio X Cell). Control mice were treated with 50 µg of rat IgG2a isotype antibody (Bio X Cell) via intraperitoneal route at 1 hour before infection. Myeloid cell depletion was performed as previously reported (11, 64). Mice received intraperitoneal injections of 250µg of either anti-GR1 (clone RB6-8C5, BioXCell) or IgG2b isotype (clone LFT-2, BioXCell) per mouse on days 3, 6, and 9 post IAV infection. The depletion efficiency was tested by flowing staining on day 7 after infection.

## COMPUTATIONAL METHODS

### Sequence alignment and identification of Virus^+^ cells

A combined mouse (mm10 transcriptome) and influenza A virus PR8 (NCBI records: NC_002016, NC_002017, NC_002018, NC_002019, NC_002020, NC_002021, NC_002022, NC_002023) reference genome was created as described previously (65) using the mkref command from the CellRanger toolkit (10 Genomics, version 7.0.1). Quantification of host and viral transcripts was performed using the CellRanger count command. To prevent the loss of cells with lower RNA counts such as neutrophils, CellRanger count was run with the following non-default parameters, as suggested by 10X Genomics: force_cells=10000 and include_introns=true. Cells were annotated Virus^+^ if expression of any of the 8 segments of the influenza genome (annotated within the genome as IAV1-IAV8) was greater than zero, and in total we identified 6,216 Virus^+^ cells (2,977 from 7 DPI vehicle and 3,239 from 7 DPI RTX). As expected, no Virus^+^ cells were detected in the two naïve samples.

### Dimensionality reduction and cell type annotation

Cells with more than 25% mitochondrial reads were filtered out during pre-processing, yielding 37,543 high quality cells. Normalization, variable feature selection (5000 genes), scaling, dimensionality reduction (PCA, top 100 components selected), and clustering were performed using Seurat. An initial round of Leiden clustering was performed using the Seurat FindClusters function at a resolution of 0.25 to identify lineages: 17,667 myeloid cells (expressing *Sirpa* and *Csf2ra*) and lymphoid (expressing *Ets1* and *Ikzf3*) 9,775 immune cell cells, and 10,101 lower quality cells and doublets. The dataset was then split into myeloid and lymphoid subsets, which were each re-scaled prior to dimensionality reduction and clustering as described above. For within-lineage Leiden clustering, we used resolution of 5, and then recursively merged highly similar clusters (Pearson’s *R*>0.8 between mean PC scores across all cells in each cluster). The resulting clusters were then annotated based on expression of known marker genes (**Table 2**). To identify cell type-specific markers from our dataset that were consistent across treatment and viral exposure, we adjusted for these covariates when performing differential expression analysis using the limma package (66). We computed all pairwise comparisons of cell type clusters, and markers were then ranked by the maximum Q-value and minimum log_50_-fold change, as described previously (67).

For detailed analysis of neutrophils, we isolated the 2,921 in the relevant cluster, and then re-scaled the data and performed variable feature selection as described above. For dimensionality reduction, the top 25 principal components were selected. We then performed Leiden clustering at a resolution of 0.25; one small cluster of 52 cells showed a higher percentage of mitochondrial reads (but less than our threshold of 25%) and was merged with its most similar (Pearson’s correlation=0.62) cluster, to result in our final 3 neutrophil clusters. Neutrophil subtype-specific marker genes were identified as described in the previous paragraph.

### Testing for changes in cell proportions in the scRNAseq dataset

To test for changes in cell type proportions, we fit a logistic regression model for each cell type using the glm package in R, modeling the probability of each cell being of that type as a binomial variable. The point estimate of this model is the log fold-change in the proportion of a given cell type between vehicle and RTX. We used the same approach to test for differences in the proportion of Virus^+^ cells.

### Identification of differentially regulated genes and cytokine signatures

For each cell type with at least ten cells per condition (naïve vehicle, naïve RTX, 7 DPI vehicle, and 7 DPI RTX), we used limma (66) to identify genes where the effect of viral exposure is significantly modified by RTX treatment(66). This approach fits a linear model for each gene using the RTX treatment and viral exposure group status as predictors, where the interaction term between treatment and viral exposure represents the change in slope between vehicle and RTX treatment; we refer to genes with a significant interaction term as *differentially regulated.* To identify enriched pathways among the differentially regulated genes, we tested for overlap with the “MSigDB Hallmark 2020” database (68) using the enrichR package and cell type-specific cytokine-induced signatures from a study in which mice were treated with individual cytokines and then immune cells were isolated and sequenced (47). We determined significant enrichment using a hypergeometric test (*p* < 0.05).

### Normalization of differentially regulated genes

We used multiple linear regression models to adjust for the effect that differences in cell type abundance and library complexity may have on statistical power to identify differentially regulated genes (for example, neutrophils have on average fewer transcripts detected, so changes detected can be larger relative differences). We first fit two linear regression models at the cell type level in which the average number of genes expressed was used to predict (1) cell abundance and (2) the number of differentially regulated genes. We then regressed out the effect of the average number of genes expressed by using the residuals of the first model to predict the residuals of the second model. This final model provides the relationship between cell type abundance and the number of differentially regulated genes after adjusting for the effect of the average number of genes expressed on each variable.

### Statistical analysis

Except for sequencing data, all statistical analysis was performed using Prism (version 10, GraphPad Software). Significance tests and the exact number of biological replicates are specified in the relevant figure legend. For microscopy analysis, images were blinded prior to scoring and quantification. Data are represented as mean ± standard error of the mean (SEM) throughout, and the level of significance is indicated by exact p-values.

## ACKNOWLEDGMENTS

We thank Eriko Kudo, Akiko Iwasaki, Bishi Fu, and Martin Dorf for help with establishing the IAV PR8 infection model. We thank Shuang Yu and Ruslan Medzhitov for support with the IAV WSN infections. We thank Samantha Choi and Chambit Paik for technical help. We thank Sarah Fortune, Susan Swain, Antonia Wallrapp, Amre Shalaby, and Wenjiang Deng for helpful discussions. We thank Mark Hoon (NIH) for *Trpv1*-DTR mice. We thank the MicRoN (Microscopy Resources on the North Quad) Core for imaging support; the HMS Immunology flow cytometry core for flow cytometry analysis; the Harvard Bauer Core Facility Sequencing team for single cell sequencing. We acknowledge funding support from National Institutes of Health (NIH) 5R01DK127257 and 5R01AI168005, Burroughs Wellcome Fund, Jackson-Wijaya Fund, and Food Allergy Science Initiative to I.M.C.; the Parker B. Francis Fellowship to A.L.H.; NIGMS R35GM155117, NIGMS P20GM113117, and American Lung Association to P.B; NIH 5T32HL007118 to N.A.; R01AI143861 to K.M.K; NIH F31 HL132645 to B.D.U.

## Author Contributions

Conceptualization: P.B., J.X., D.Y., N.A., S.U., and I.M.C.; Methodology: A.L.H., P.B., D.Y., N.A., J.X., B.D.U., and S.U.; Investigation: P.B., J.X., D.Y., N.A., S.U., S.T.Y., D.A.H., B.D.U., O.E., N.S., A.B-D., N.B., K.A.M., R.A.F., and J.L.; Writing – original draft: N.A., D.Y., and P.B.; Writing – Editing and Finalization: A.L.H., S.U., R.A.F., K.M.K., and I.M.C.; Resources and Funding: K.M.K, B.G.Y., S.D.L., A.L.H. and I.M.C.

## Competing Interests

I.M.C. consults for GSK pharmaceuticals, Panther Life Sciences, Fzata, and Nilo Therapeutics. His lab has received sponsored research support from Abbvie and Moderna, Inc. S.U. is currently an employee of Vertex Pharmaceuticals and may own stock in the company.

## Data and materials availability

All original raw and analyzed sequencing data is available on GEO (accession GSEXXX). Multiplex cytokine array normalized data values, and the complete results of all differential expression tests from the sequencing data are provided in supplemental materials (**Tables S1-S6**). Only standard algorithms and no customized code were used to perform computational analysis. Standard R code to reproduce the main components of the analysis will be made available for research purposes upon reasonable requests to the corresponding authors.

## SUPPLEMENTARY MATERIALS

### Supplementary Figures

**Figure S1.**
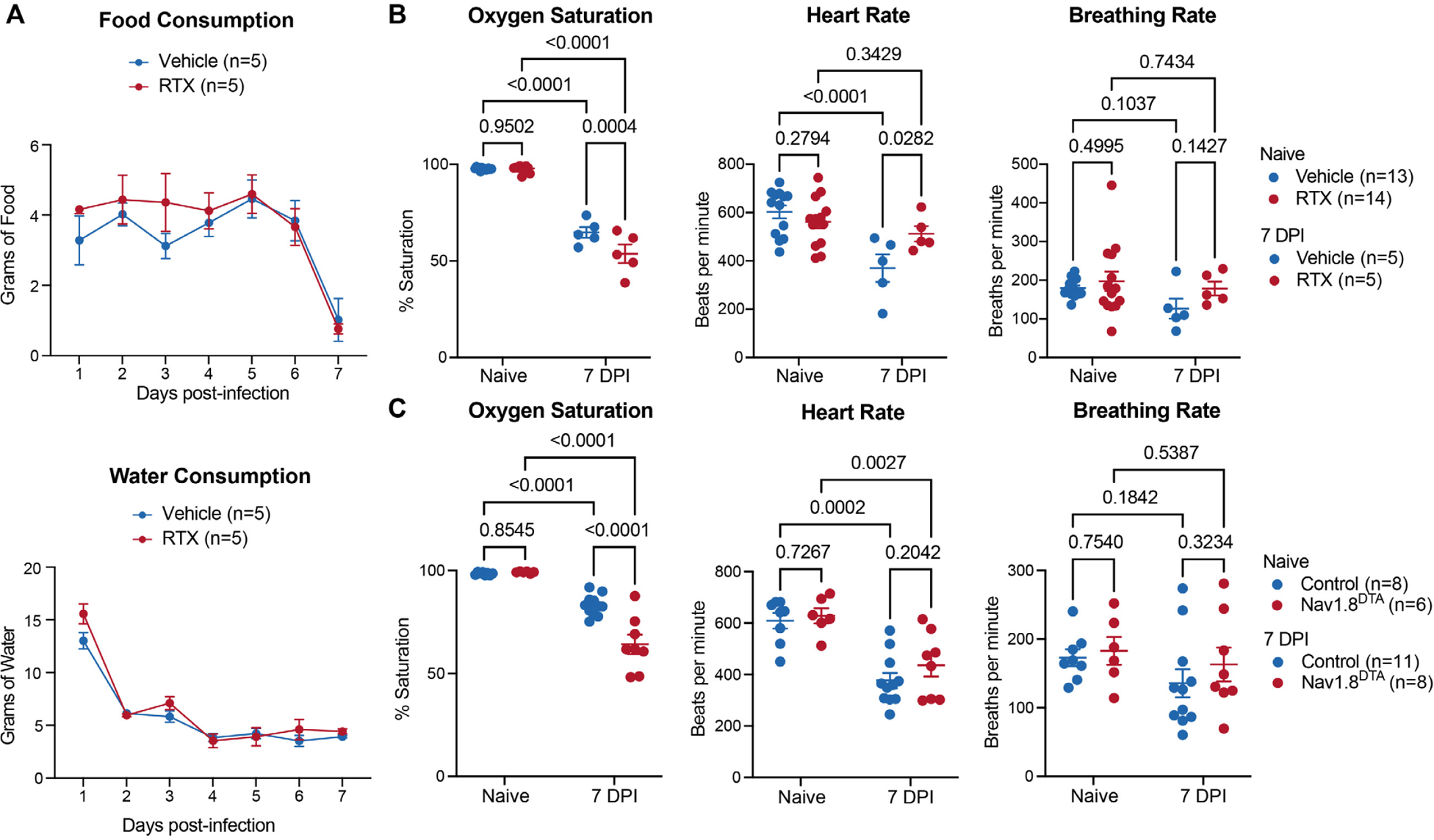
Effects of nociceptor ablation on sickness behavior and vital signs. **(A)** Daily food (top) and water (right) consumption by vehicle and RTX treated mice were monitored post-influenza infection. (n=5 per group). **(B)** Oxygen saturation, heart rate, and breathing rate were measured in RTX- and vehicle-treated mice at baseline (n=13-14 per group) and at 7 days post IAV PR8 infection (n=5 per group). **(C)** Oxygen saturation, heart rate, and breathing rate were measured in Nav1.8^DTA^ mice and littermate controls at baseline (n=6-8 per group) and at 7 DPI (n=8-11 per group). Multiple t test in (A); Two-way ANOVA in (B and C); error bars show mean±SEM; P values labeled.

**Figure S2.**
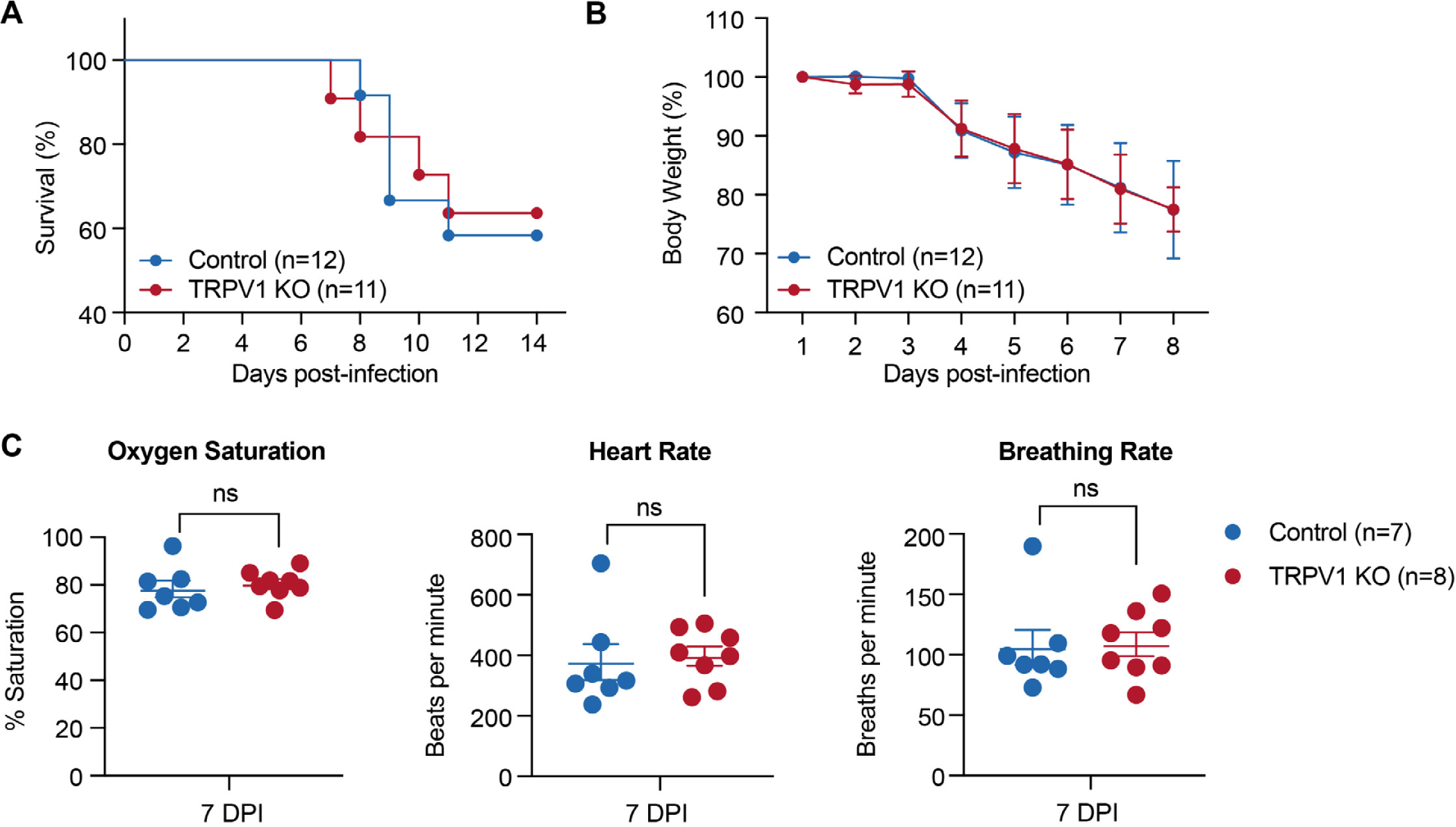
TRPV1 ion channel does not mediate influenza-induced mortality and physiological changes. **(A)** The survival curve and **(B)** bodyweight loss of *Trpv1* knockout and littermate control mice at the indicated time points post infection with 50 TCID_50_ PR8 (n=11-12 per group) **(C)** Oxygen saturation, heart rate, and breathing rate were measured in the *Trpv1* knockout and littermate control mice 7 days post IAV infection. (n=7-8 per group). Log-rank test was used in (A); Multiple t test in (B); Student’s t test in (C); error bars show mean±SEM; P values labeled.

**Figure S3.**
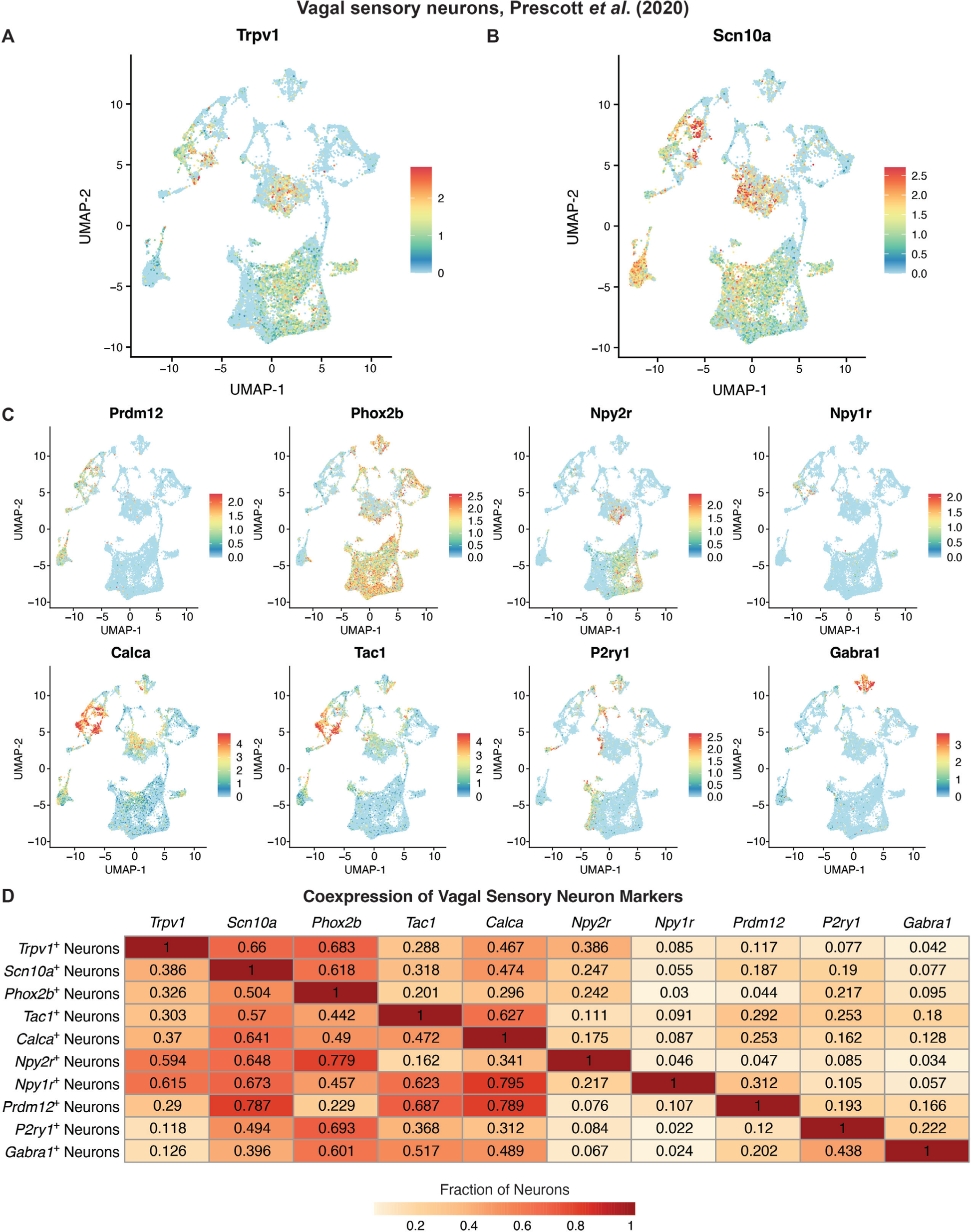
Vagal sensory neuron expression of TRPV1 and subtype-specific markers. **(A-B)** UMAP of published single-cell RNA sequencing dataset of vagal sensory neurons (Prescott *et al*. 2020) showing expression in log_50_(CPM+1) (color legend) of nociceptor markers *Trpv1* (A) and *Scn10a* (B). **(C)** UMAP displaying expression in log_50_(CPM+1) (color legend) of indicated markers of vagal sensory neuron subtypes. **(D)** Table showing the fraction of cells (color legend) from the indicated vagal sensory neuron populations (rows) that are expressing (log_50_(CPM+1) > 0) each of the neuron subset markers (columns).

**Figure S4.**
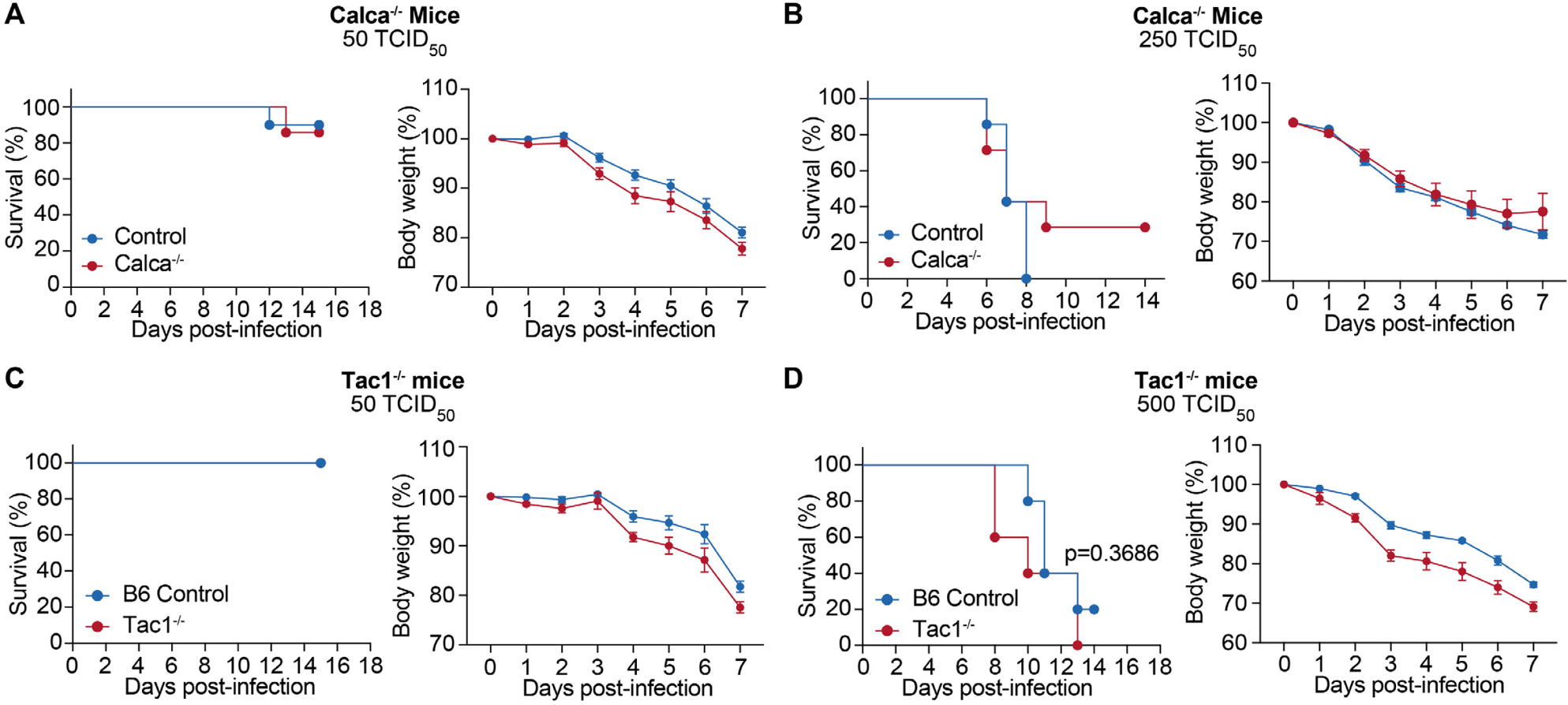
Neuropeptides CGRP and Substance P do not regulate survival in influenza infection. **(A)** The survival curve (left) and bodyweight loss (right) of *Calca* knockout and littermate control mice at the indicated time points post IAV PR8 infection with a dose of 50 TCID_50_ per mouse (n=7-10 per group). **(B)** The survival curve (left) and bodyweight loss (right) of *Calca* knockout and littermate control mice at the indicated time points post IAV PR8 infection with a dose of 250 TCID_50_ per mouse (n=7-8 per group). **(C)** The survival curve (left) and bodyweight loss (right) of *Tac1* knockout and littermate control mice at the indicated time points post IAV PR8 infection with a dose of 50 TCID_50_ per mouse (n=4-6 per group). **(D)** The survival curve (left) and bodyweight loss (right) of Tac1 knockout and littermate control mice at the indicated time points post IAV PR8 infection under a dose of 500 TCID_50_ per mouse (n=5 per group). Log-rank test for survival analysis; Multiple t test for analysis of bodyweight loss; error bars show mean±SEM; P values labeled.

**Figure S5.**
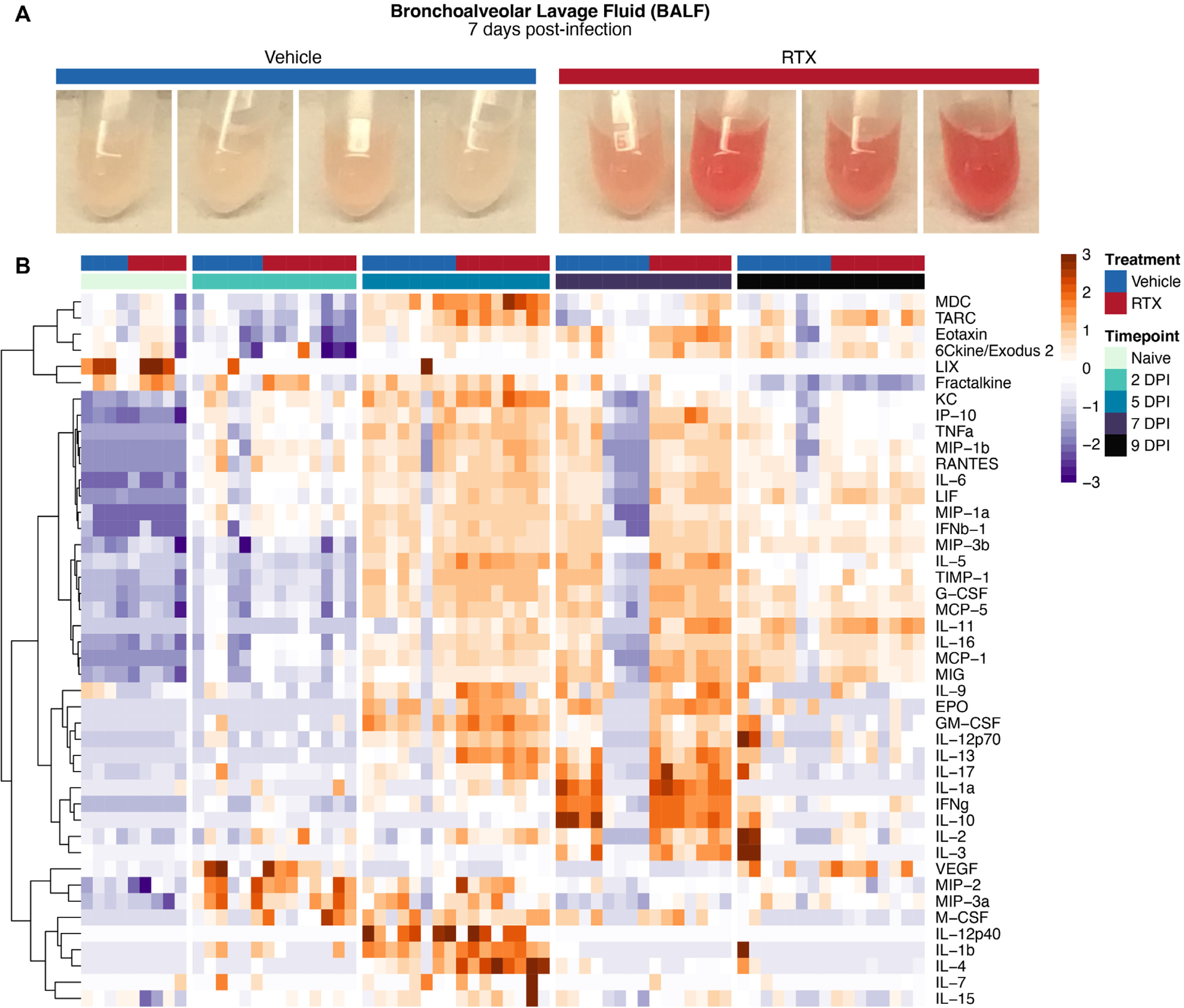
Effects of nociceptor ablation on cytokine induction during influenza infection. **(A)** Images of bronchoalveolar lavage fluid collected from vehicle (left) and RTX-treated (right) mice at day 7 post-infection. **(B)** Heatmap of protein levels (row-wise z-score of log_50_(pg/ml + 1), color bar) of the indicated cytokines and chemokines detected by the 44-Plex Discovery Assay® Array in the BALF collected on days 0 (Naïve), 2, 5, 7, and 9 post IAV PR8 infection. (n=4-8 per group).

**Figure S6.**
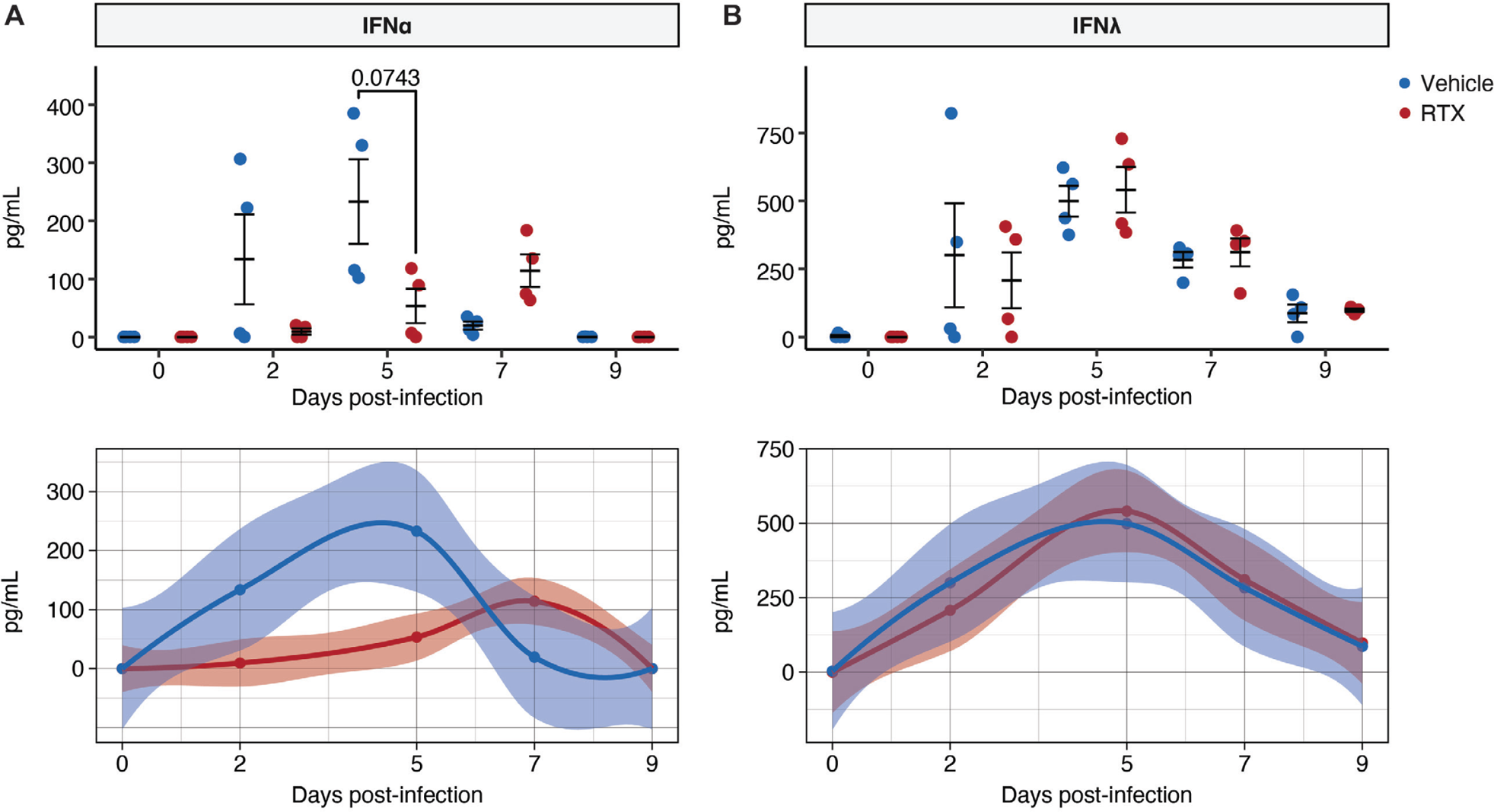
Effect of nociceptor ablation on interferon induction after influenza infection. **(A-B)** Protein levels (in pg/mL, y axis) (top) and LOESS regression curve of protein levels (bottom) detected for **(A)** IFNα and **(B)** IFNλ measured in BALF by ELISA at indicated time points post-infection (x axis). (n=4 per group). Two-way ANOVA with FDR adjusted p values in (B-C); error bars show mean±SEM.

**Figure S7.**
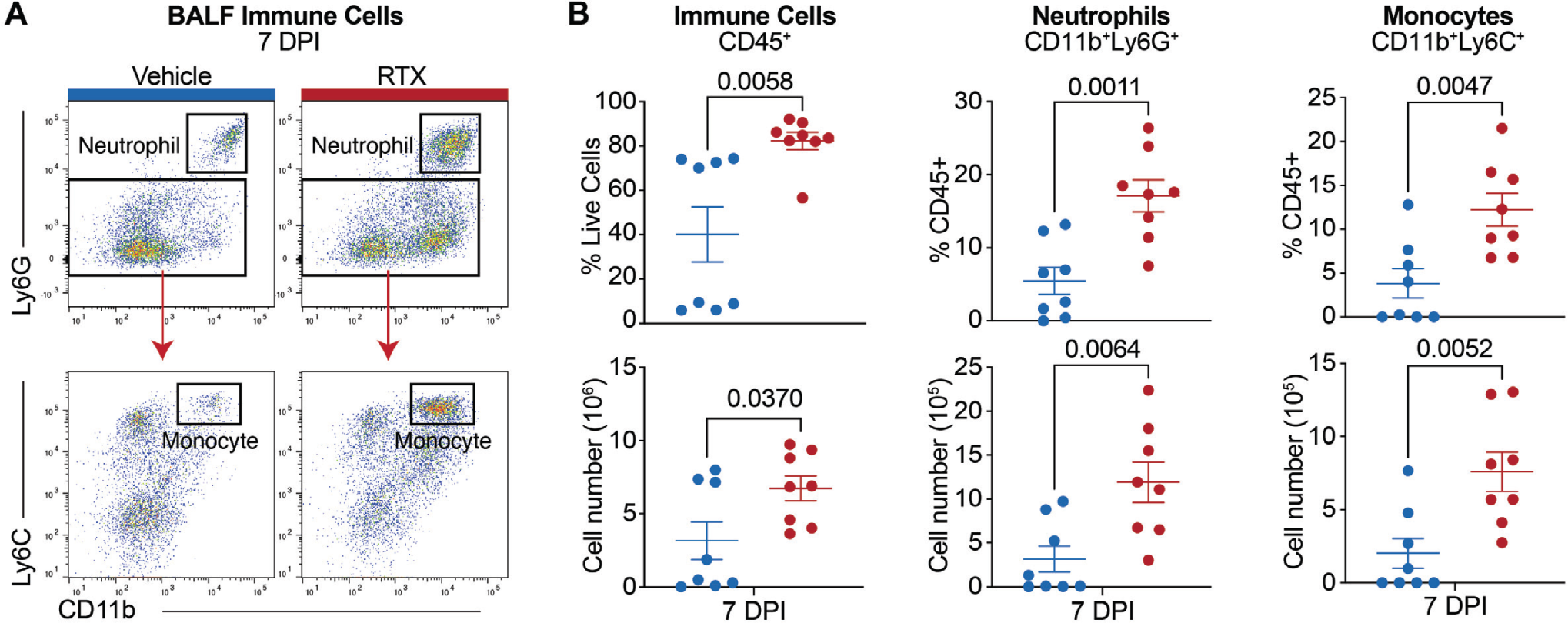
Flow cytometry of immune cells in the bronchoalveolar lavage fluid. **(A)** Representative flow plots of live CD45^+^ cells isolated from BALF of vehicle (left) and RTX-treated mice at 7 DPI, gated for neutrophils (Ly6G^+^CD11B^+^) and monocytes (Ly6G^−^ Ly6C^+^CD11B^+^). **(B)** Quantification of BALF CD45^+^ cells, neutrophils, and monocytes as a percentage (top) or absolute cell numbers (bottom). (n=8 per group). Student’s t test in (B); error bars show mean±SEM; P values labeled.

**Figure S8.**
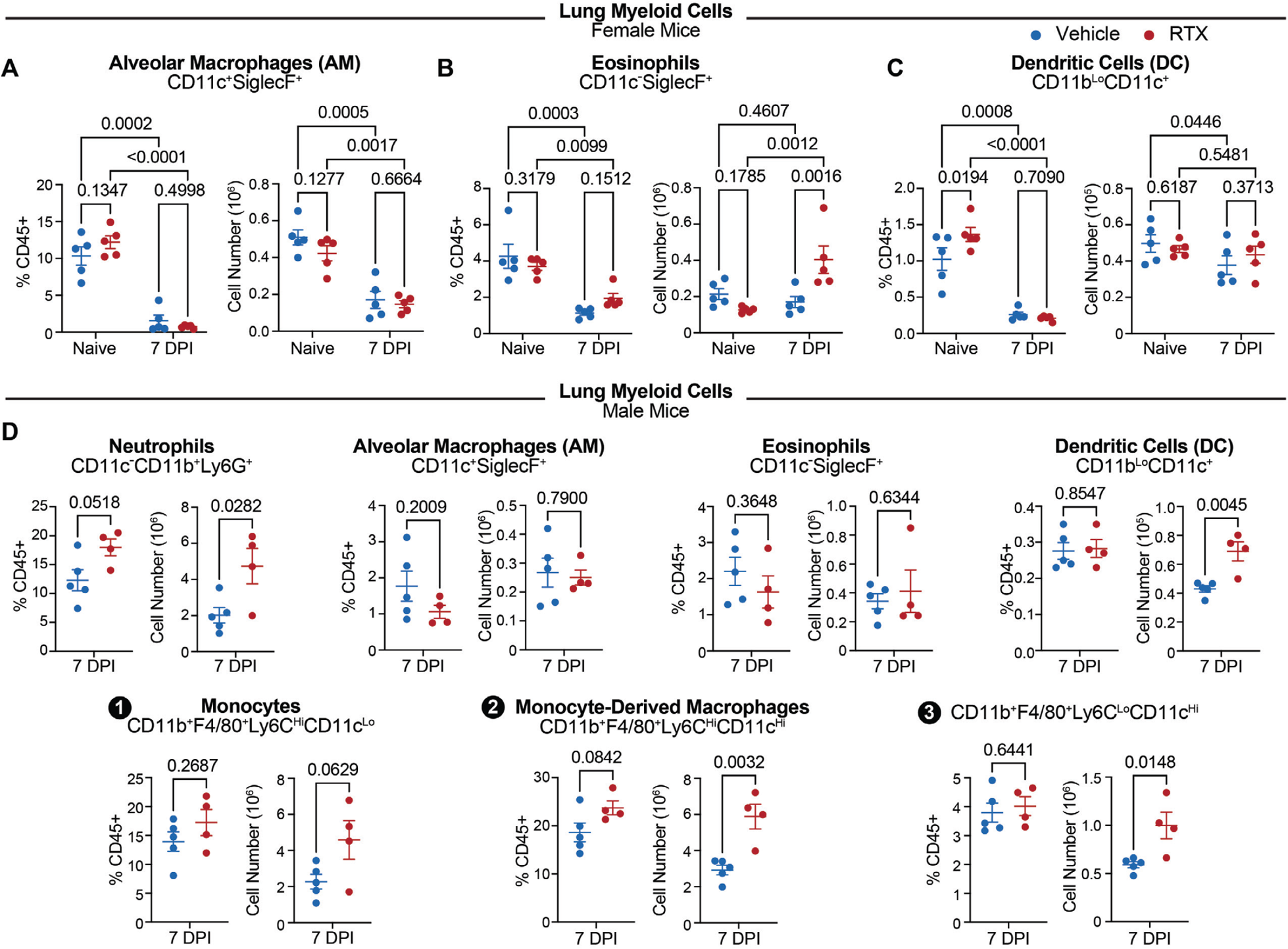
Changes in lung myeloid immune cell populations after influenza infection. **(A-C)** Quantification of labeled myeloid populations in the lung tissue of vehicle- and RTX-treated female mice at baseline and on day 7 post IAV PR8 infection shown as a percent of CD45^+^ immune cells (left) and absolute cell numbers (right). (n=5 per group). **(D)** Quantification of indicated myeloid populations in the lung tissue of vehicle and RTX-treated male mice at day 7 post-influenza infection shown as a fraction of CD45^+^ immune cells (left) and absolute cell numbers (right). (n=4-5 per group). Two-way ANOVA in (A-C); Student’s t test in (D); error bars show mean±SEM; P values labeled.

**Figure S9.**
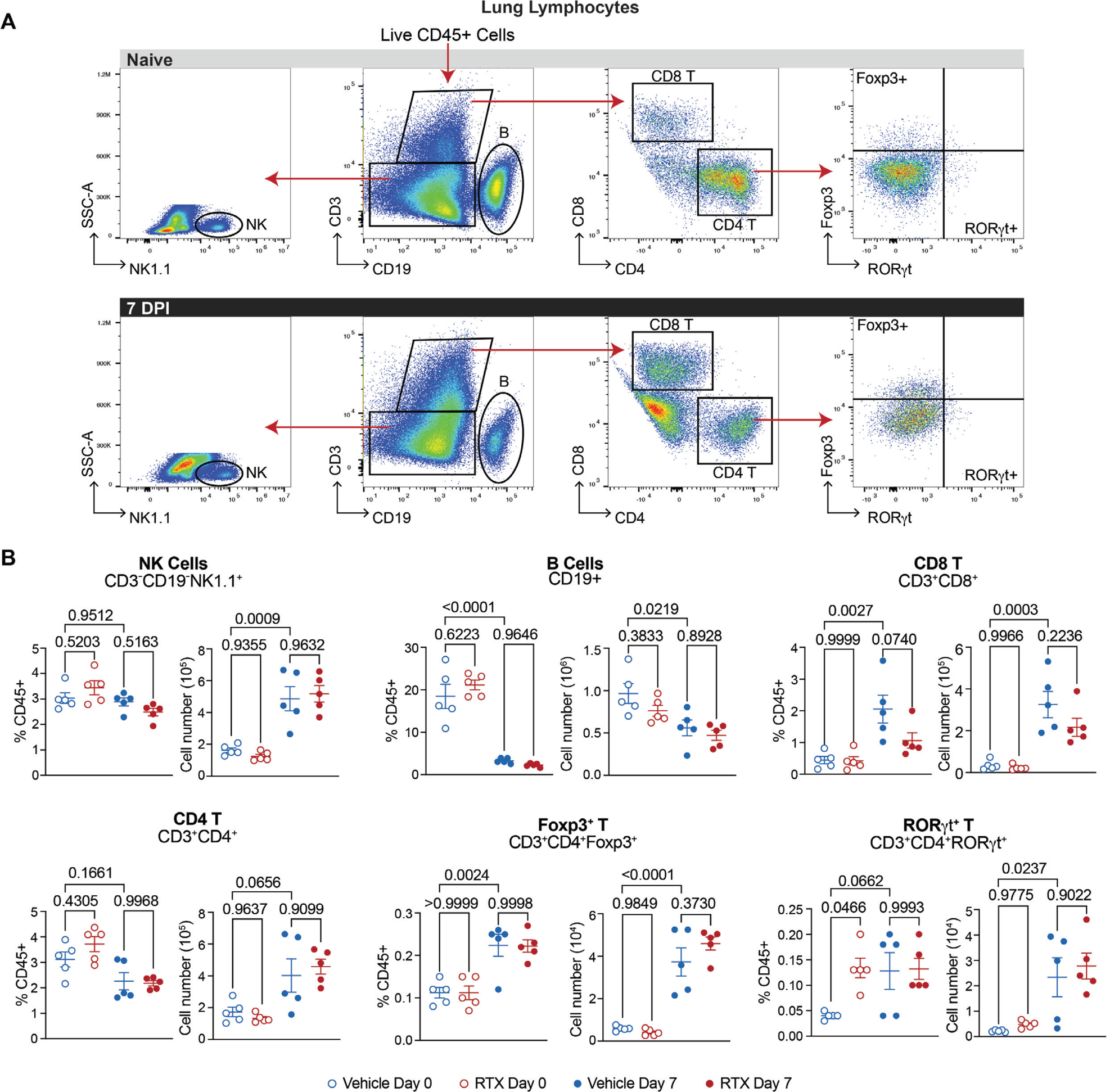
Changes in lung lymphoid immune cell populations after influenza infection. **(A)** Representative flow plots and gating strategy for the indicated lymphoid cell populations isolated from mouse lung tissue of naïve mice (top) and at 7 days post-infection (bottom). **(B)** Quantification of labeled lymphoid populations in the lung tissue of vehicle and RTX-treated mice at baseline and on day 7 post IAV PR8 infection shown as a fraction of CD45^+^ immune cells (left) and absolute cell numbers (right). (n=5 per group). One-way ANOVA in (B); error bars show mean±SEM; P values labeled.

**Figure S10.**
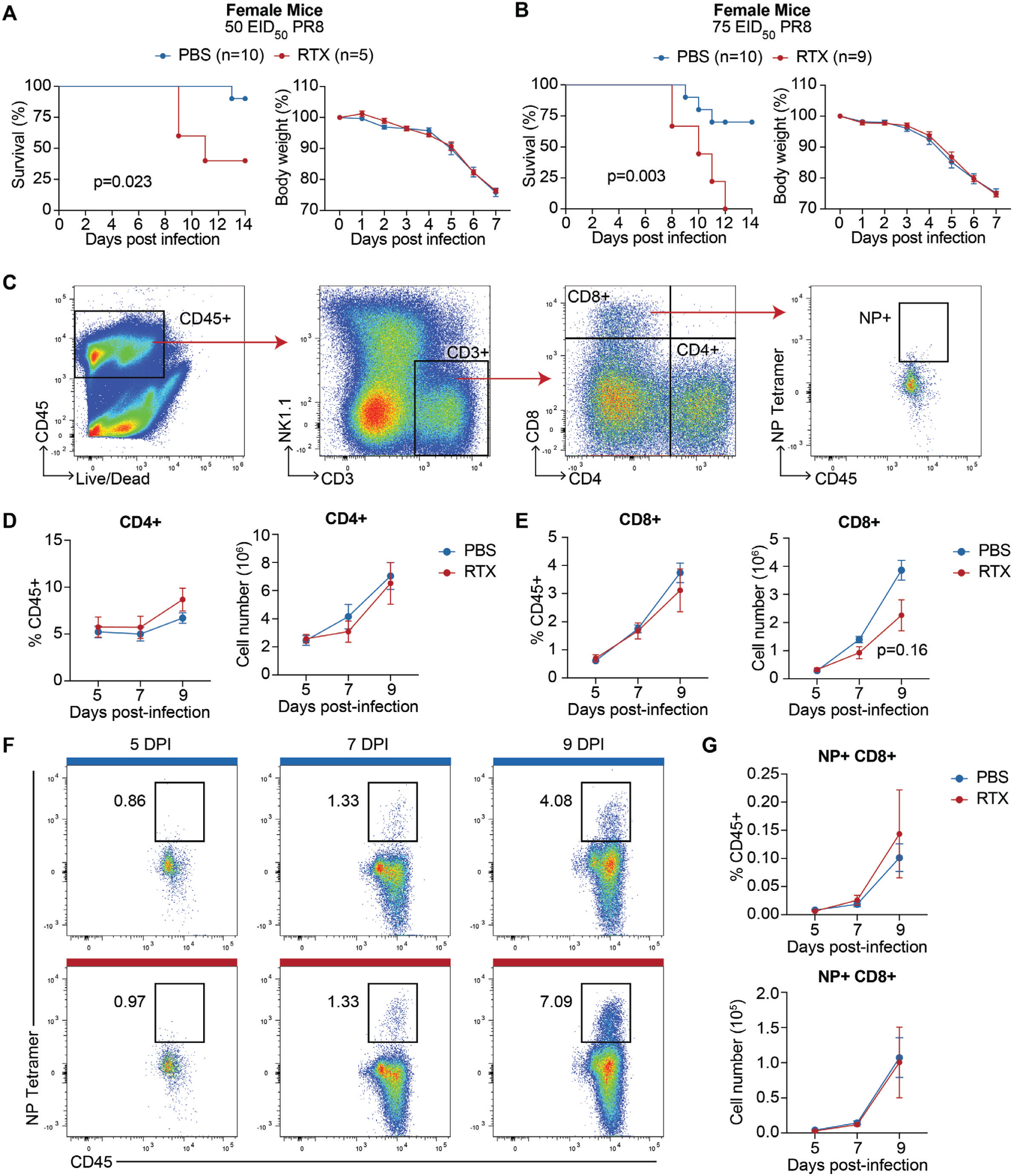
Nociceptor deficient exhibit no difference in influenza virus specific CD8 T cells. **(A-B)** Female WT mice (at NYU) were treated with RTX or PBS control then intranasally infected with IAV PR8 at a dose of **(A)** 50 EID_50_ or **(B)** 75 EID_50_ four weeks later and monitored for survival (left) and changes in body weight (right). (n=5-10 per group). **(C)** Representative flow plots and gating strategy for influenza virus-specific (IAV NP tetramer^+^) CD8 T cells after infection with IAV PR8 at a dose of 75 EID_50_. **(D)** Quantification of CD4^+^ T cells in the lung tissue of RTX and PBS-treated mice at indicated timepoints post IAV PR8 (75 EID_50_) infection shown as a percentage of CD45^+^ immune cells and absolute cell numbers. (n=4 per group). **(E)** Quantification of CD8^+^ T cells in the lung tissue of PBS and RTX-treated mice at indicated timepoints post IAV PR8 (75 EID_50_) infection shown as a percentage of CD45^+^ immune cells and absolute cell numbers. (n=4 per group). **(F)** Representative flow plots of CD8 T cell staining for IAV nucleoprotein tetramer at indicated time points post IAV PR8 (75 EID_50_) infection. **(G)** Quantification of influenza virus specific (IAV NP tetramer^+^) CD8 T cells in the lung tissue of PBS and RTX-treated mice at indicated timepoints post-infection shown as a percentage of CD45+ cells and absolute cell numbers. (n=4 per group). Log-rank test was used for survival analysis; Multiple t test for body weight analysis; Two-way ANOVA in (D-E, and G); error bars show mean±SEM; P values labeled.

**Figure S11.**
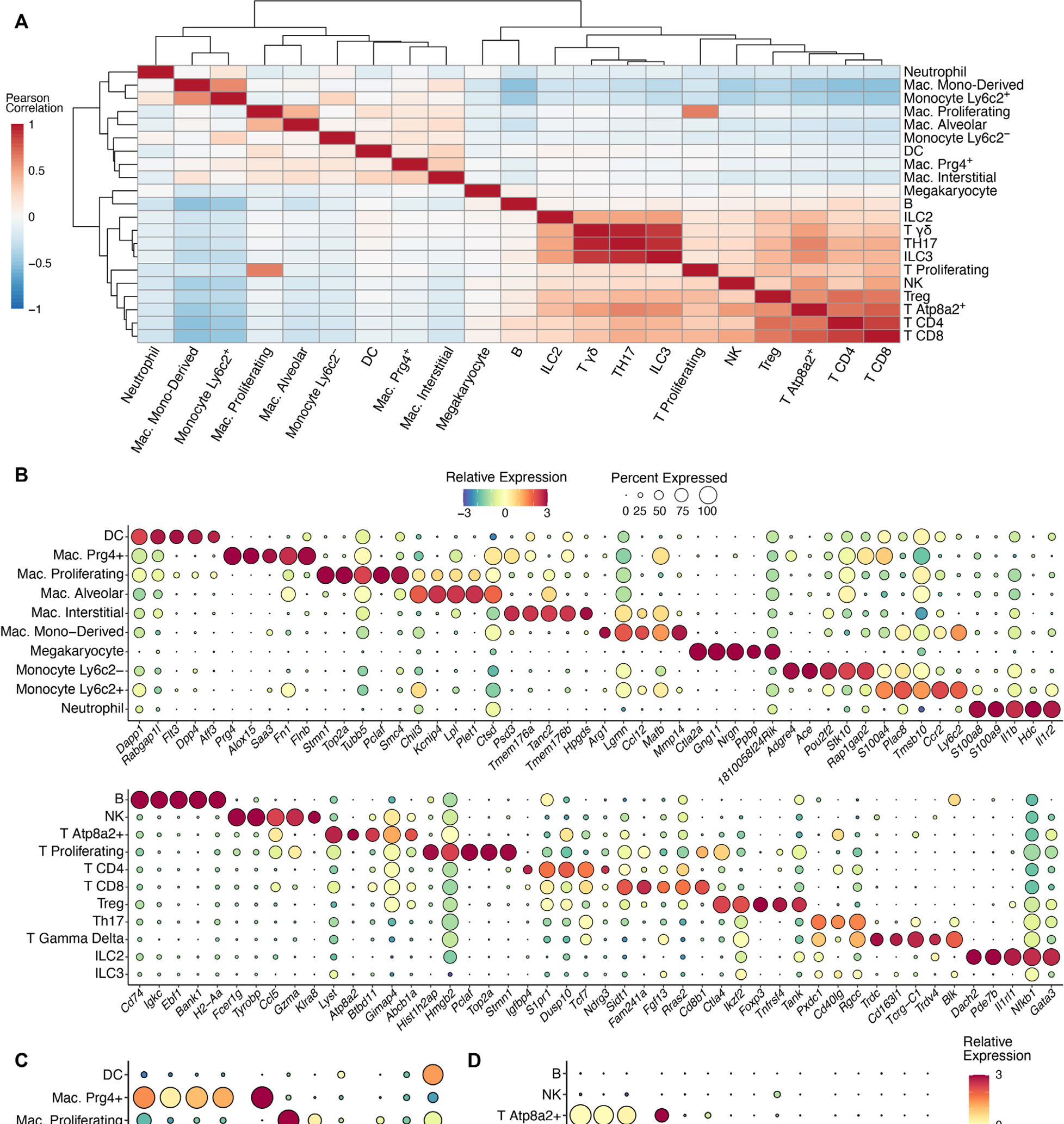
Identification of lung immune cell populations by single-cell RNA sequencing. **(A)** Hierarchically clustered heatmap showing the similarity (Pearson correlation coefficient, color legend) between identified immune cell types. **(B)** Dot plots showing the top cell type specific marker genes (x axis) for each myeloid (top) and lymphoid (bottom) cell type (y axis). Color of the dot indicates the scaled average expression (z-score of log_50_(CPM+1), color bar) of a gene within a cluster and the size of the dot corresponds to the fraction of cells in a cluster expressing the gene. **(C)** Dot plot showing expression within myeloid cells of shared macrophage genes, macrophage subtype specific genes, and relevant genes from the literature, visualized as in (B). **(D)** Dot plot showing expression within lymphoid cells of shared T cell genes, T cell subtype-specific genes, and shared T_50_17/ILC3 genes, visualized as in (B-C).

**Figure S12.**
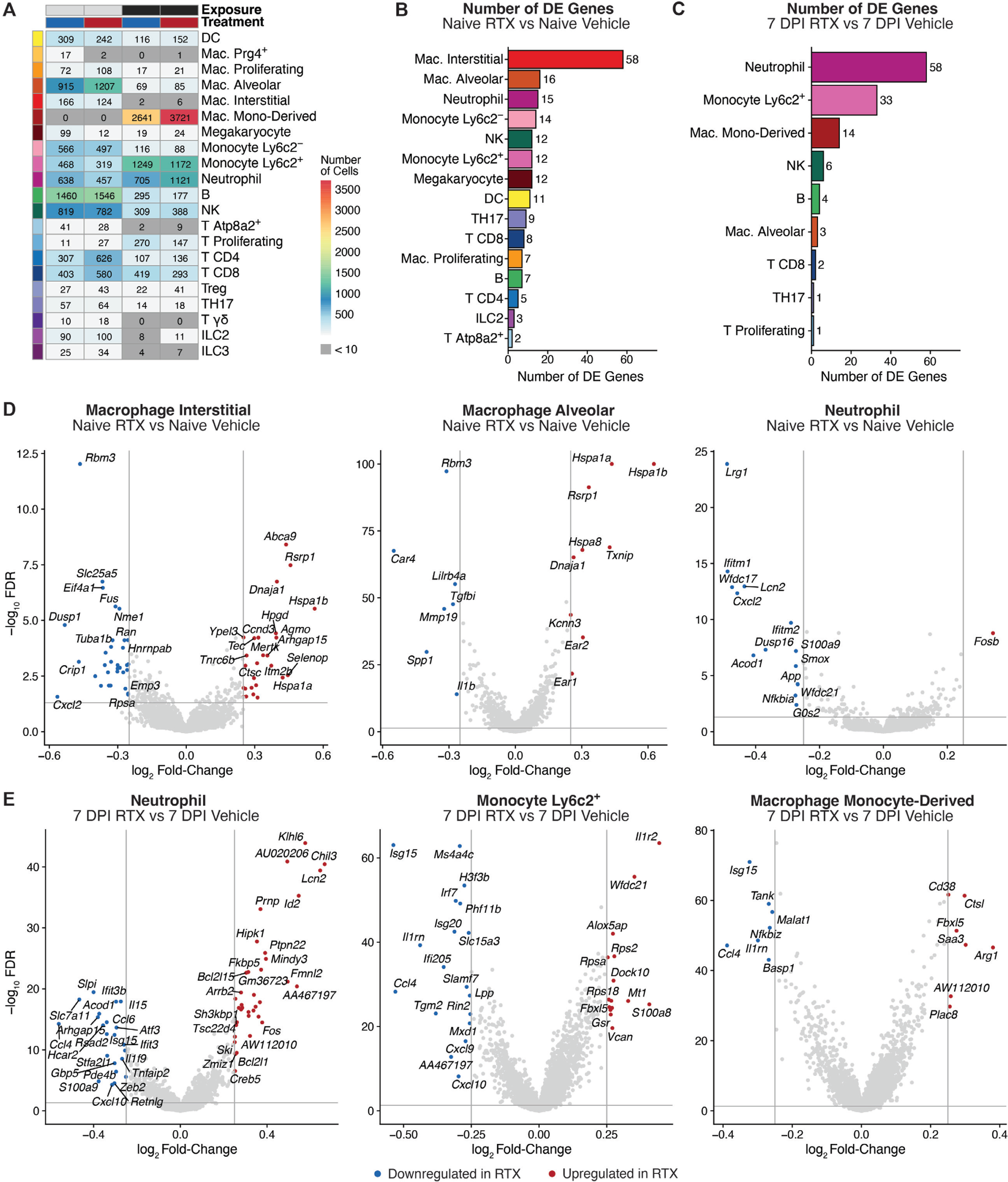
Nociceptor ablation alters the transcriptomes of lung immune cell types at baseline and after influenza infection. **(A)** Table indicating the number of cells (label, color legend) of each immune cell type from the indicated treatment and exposure groups (color bars on top) in the single-cell RNA sequencing dataset. **(B)** Bar plot indicating the number of differentially expressed genes (x axis, label) identified for each cell type (y axis) between RTX and Vehicle at baseline. **(C)** Bar plot indicating the number of differentially expressed genes identified for each cell type between RTX and Vehicle at 7 days post-infection, visualized as in (B). **(D)** Volcano plots for the indicated cell types illustrating the log_50_ fold-change (x axis) and −log_50_ FDR-adjusted p value (y axis) of each gene when comparing the cells from Naïve RTX and Naïve Vehicle. Horizontal and vertical lines illustrate the significance thresholds, and points corresponding to a differentially expressed gene significantly upregulated in RTX (positive log_50_ FC) are colored red and genes significantly downregulated (negative log_50_ FC) are colored blue. **(E)** Volcano plots for the indicated cell types illustrating the log_50_ fold-change (x axis) and −log_50_ FDR-adjusted p value (y axis) of each gene when comparing the cells from 7 DPI RTX and 7 DPI Vehicle, visualized as in (D).

**Figure S13.**
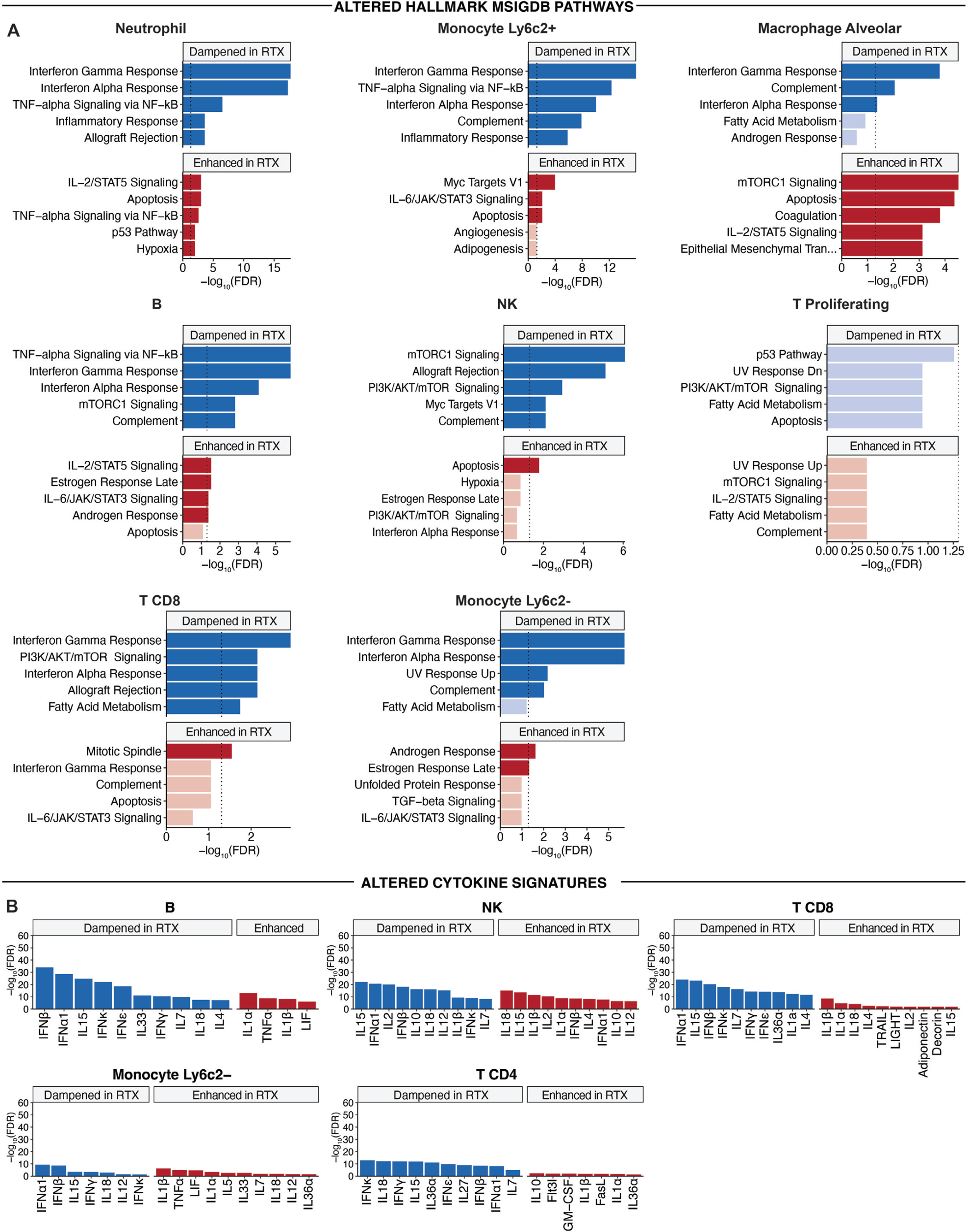
Genes differentially regulated between vehicle and RTX exhibit altered pathway and cytokine signatures. **(A)** Bar plots showing the top 5 pathways (y axis) enriched among the genes dampened (top, blue bars) or enhanced (bottom, red bars) in RTX compared to Vehicle, shown for the indicated cell types. Vertical line illustrates the significance threshold for the −log_50_ FDR-adjusted hypergeometric p value (x axis). Darker-colored bars correspond to significant pathways. **(B)** Bar plot indicates the −log_50_ FDR-adjusted hypergeometric p value (y axis) for the overlap of the genes dampened (left, blue bars) or enhanced (right, red bars) in RTX with cell-type specific gene signatures of individual cytokines (x axis).

**Figure S14.**
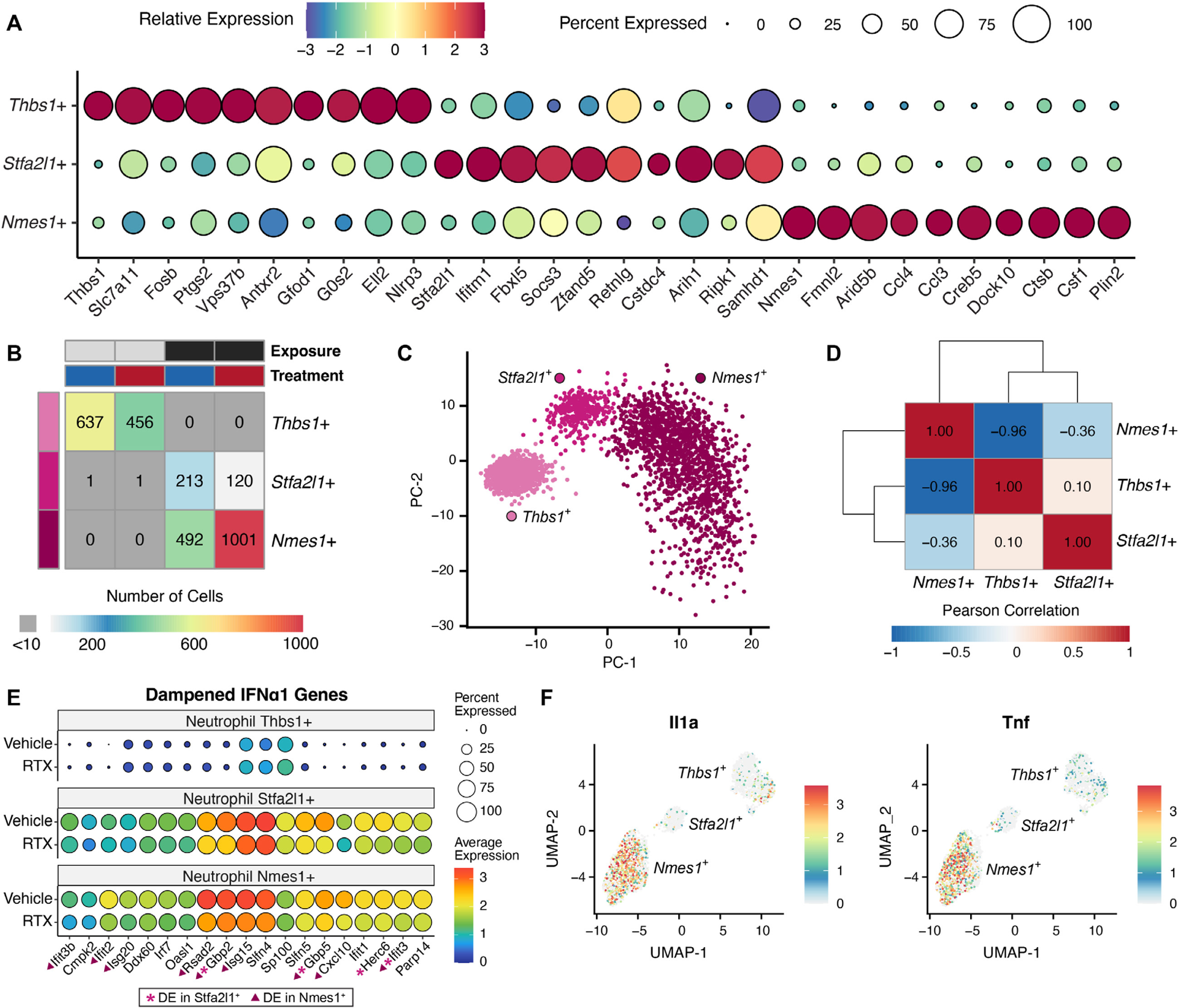
Identification of transcriptionally distinct neutrophil subsets. **(A)** Dot plot showing the top cluster-specific marker genes (x axis) for each neutrophil subset (y axis). Color of the dot indicates the scaled average expression (z-score of log_50_(CPM+1), color bar) of a gene within a cluster and the size of the dot corresponds to the fraction of cells in a cluster expressing the gene. **(B)** Table indicating the number of cells (label, color legend) of each neutrophil subset isolated from the indicated treatment and exposure groups (color bars on top). **(C)** PCA plot showing separation of the three identified neutrophil subsets (color and label) by the first two principal components. **(D)** Hierarchically clustered heatmap showing the similarity (Pearson correlation coefficient, color legend and labels) between cells from each neutrophil subset. **(E)** Dot plot showing the expression within specific neutrophil subsets of IFNα1 signature genes dampened in RTX neutrophils. Color of the dot indicated the average expression (log_50_(CPM+1), color bar) of a gene within a cluster and the size of the dot corresponds to the fraction of cells in a cluster expressing the gene. **(F)** UMAP plots showing the expression (log_50_(CPM+1), color bar) of *Il1a* and *Tnf* by neutrophils.

**Figure S15.**
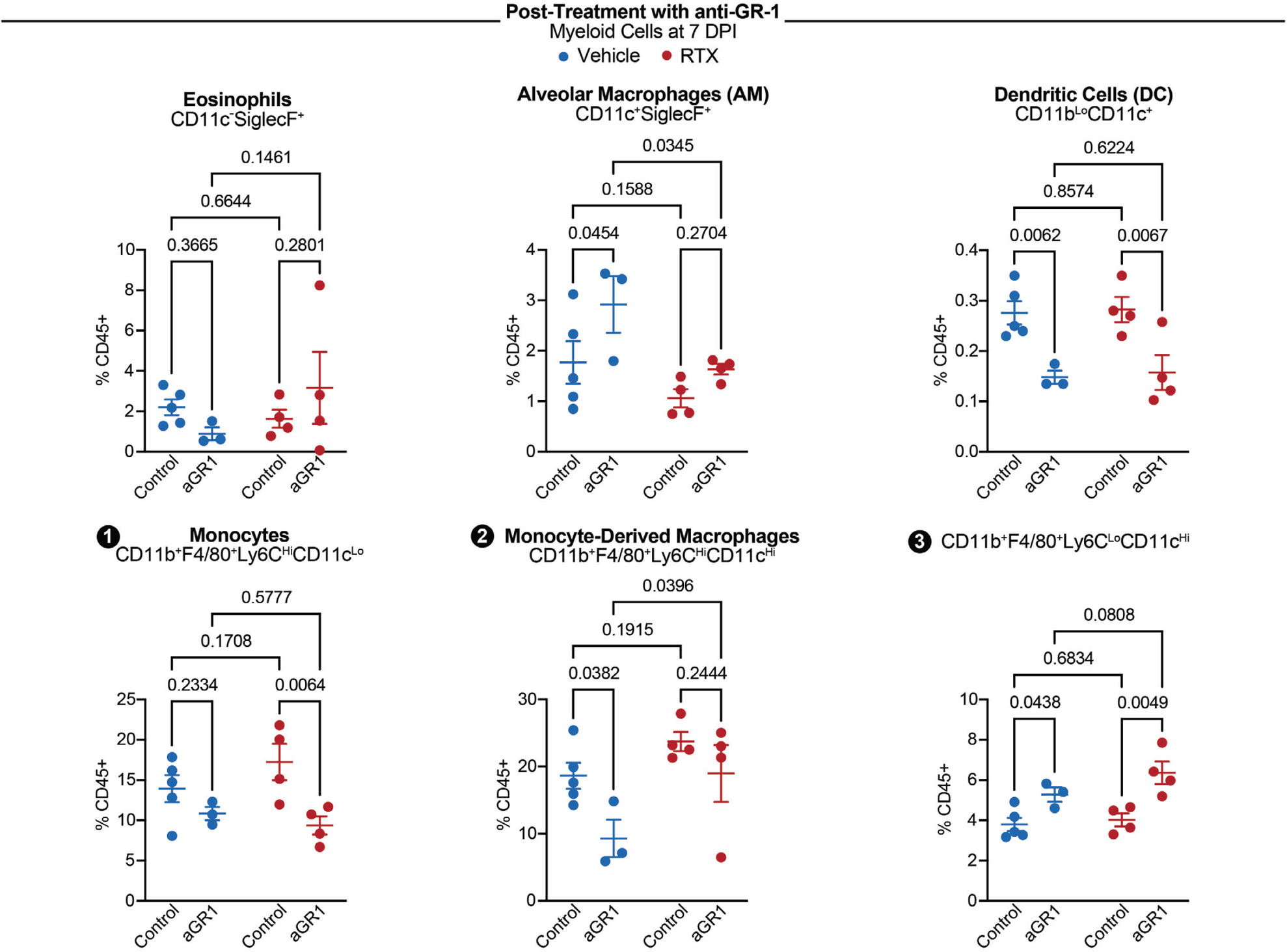
Effect of post-treatment with anti-GR-1 on lung myeloid cell populations. Quantification of labeled myeloid populations at day 7 post IAV PR8 infection in the lung tissue of vehicle and RTX-treated control mice and mice treated with anti-GR-1 (250ug on days 3 and 6 post-infection), shown as a percent of CD45^+^ immune cells (left) and absolute cell numbers (right). (n=3-5 per group). Two-way ANOVA; error bars show mean±SEM; P values labeled.

### Supplementary Tables

**Table S1. Cytokine protein levels in the bronchoalveolar lavage fluid.** Cytokine protein levels (in pg/mL) detected by the 44-Plex Discovery Assay® Array of the bronchoalveolar lavage fluid from vehicle and RTX-treated mice at days 0 (naïve), 2, 5, 7, and 9 post-infection with IAV PR8.

**Table S2. Literature-derived lung immune cell type marker genes.** Immune cell type marker genes from the literature that were used to annotate clusters in the single-cell RNA sequencing dataset.

**Table S3. Significant marker genes for lung myeloid and lymphoid immune cell types.** Significant (maximum FDR < 0.05 and minimum log_50_ fold-change > 0.25) marker genes for each cell type as identified by pairwise differential expression tests performed with *limma*, separated into myeloid and lymphoid immune compartments and ranked by minimum fold-change.

**Table S4. Molecular signatures of lung immune cell types following nociceptor ablation and viral exposure.** Significant (FDR < 0.05 & absolute value of log_50_ fold-change > 0.25) differential expression results from (1) head-to-head comparisons between RTX and Vehicle, in Naïve or 7 DPI viral exposure groups and (2) genes where the effect of viral exposure is significantly modified by RTX treatment (differentially regulated genes).

**Table S5. Enriched pathways and cytokines among differentially regulated genes.** Pathways and cytokine signatures enriched (*p* < 0.05, hypergeometric test) among the genes dampened or enhanced by RTX treatment for each cell type.

**Table S6. Significant marker genes for neutrophil clusters.** Significant (maximum FDR < 0.05 and minimum log_50_ fold-change > 0.25) marker genes for each cell type as identified by pairwise differential expression tests performed with *limma*, ranked by minimum fold-change.

### Supplementary Data Files

**Data File S1**. Complete results from pairwise comparisons (Table S3) for myeloid cells

**Data File S2**. Complete results from pairwise comparisons (Table S3) for lymphoid cells

**Data File S3**. Complete differential expression results for comparisons between vehicle and RTX at baseline and at 7 DPI (Table S4)

**Data File S4**. Complete results for comparison between viral exposure groups and modified by RTX treatment (differentially regulated genes) (Table S4)

**Data File S5**. Complete results from pairwise comparisons for neutrophil clusters (Table S6)

